# Granular component sub-phases direct ribosome biogenesis in the nucleolus

**DOI:** 10.1101/2025.03.01.640913

**Authors:** Priyanka Dogra, Mylene C. Ferrolino, Suparna Khatun, Michele Tolbert, Qi Miao, Shondra M. Pruett-Miller, Aaron Pitre, Swarnendu Tripathi, George E. Campbell, Richa Bajpai, Toler Freyaldenhoven, Eric Gibbs, Cheon-Gil Park, Richard W. Kriwacki

**Affiliations:** Department of Structural Biology, St. Jude Children’s Research Hospital, Memphis, Tennessee, USA; Center for Advanced Genome Engineering, St. Jude Children’s Research Hospital, Memphis, Tennessee, USA; Department of Cell and Molecular Imaging, St. Jude Children’s Research Hospital, Memphis, Tennessee, USA; Cell and Tissue Imaging Shared Resource, St. Jude Children’s Research Hospital, Memphis, Tennessee, USA; Department of Microbiology, Immunology and Biochemistry, University of Tennessee Health Sciences Center, Memphis, Tennessee, USA

**Author notes:** Contributed equally (Priyanka Dogra, Mylene C. Ferrolino). Corresponding author: Richard W. Kriwacki. ElevateBio, 200 Smith Street, Waltham, MA 02451.

## Abstract

The hierarchical, multiphase organization of the nucleolus underlies ribosome biogenesis. Ribonucleoprotein particles that regulate ribosomal subunit assembly are heterogeneously disposed in the granular component (GC) of the nucleolus. However, the molecular origins of the GC’s spatial heterogeneity and its association with ribosomal subunit assembly remain poorly understood. Here, using super-resolution microscopy, we uncover that key GC biomolecules, including nucleophosmin (NPM1), surfeit locus protein 6 (SURF6), and ribosomal RNA (rRNA), are heterogeneously localized within sub-phases in the GC. In vitro reconstitution showed that these GC biomolecules form multiphase condensates with SURF6/rRNA-rich core and NPM1-rich shell, providing a mechanistic basis for GC’s spatial heterogeneity. SURF6’s association with rRNA is weakened upon ribosome subunit assembly, enabling NPM1 to extract assembled subunits from condensates—suggesting an assembly-line-like mechanism of subunit efflux from the GC. Our results establish a framework for understanding the heterogeneous structure of the GC and reveal how its distinct sub-phases facilitate ribosome subunit assembly.

## Introduction

The discovery of phase-separated biomolecular condensates represents a paradigm shift in our understanding of how biomolecules are organized within cells to perform their biological functions ^1–4^. Biomolecular condensates are dynamic, membraneless assemblies in cells that form through weak, transient multivalent interactions between proteins, nucleic acids, and other biomolecules ^4–6^. The nucleolus, a dynamic multiphase condensate, facilitates ribosome biogenesis through complex molecular processes supporting the assembly of pre-ribosomal particles ^7–12^. These processes are coordinated through the hierarchical spatial organization of the nucleolus. Human nucleoli contain three co-existing immiscible sub-compartments, termed the fibrillar center (FC), dense fibrillar component (DFC), and granular component (GC) ^10^. The immiscibility and spatial separation of these nucleolar sub-compartments stem from differences in their material properties, particularly surface tension ^13^. Within the FC, RNA polymerase I, aided by associated transcription factors, binds to regulatory elements within tandemly repeated ribosomal RNA (rRNA) genes (ribosomal DNA, or rDNA) and initiates transcription ^14^. Localized transcription generates nascent pre-rRNAs, which undergo extensive post-transcriptional modifications and processing within the DFC ^14^. During this stage, modifications including methylation, pseudouridylation, and cleavage events transform pre-rRNA into its mature forms ^7,10^. Notably, more than 200 assembly factors facilitate cleavage, modification, and processing of pre-rRNA to yield mature rRNAs ^15^. Following these steps, the processed pre-rRNA moves into the GC and assembles with ribosomal proteins to form ribosomal subunits ^16^. The separation of the FC, DFC, and GC phases within the nucleolus enables the compartmentalization of the different processes underlying ribosome biogenesis.

The liquid-like nature of the different sub-compartments of the nucleolus enables ribosomal components and supporting biomolecules to move within and between them in the process of ribosome subunit assembly ^13,17,18^. Numerous nucleolar proteins contain intrinsically disordered regions (IDRs) that mediate phase separation, which establishes the distinct sub-compartments within the nucleolus. Nucleolar proteins, such as nucleophosmin (NPM1), nucleolin (NCL), and fibrillarin (FBL), have been shown to undergo phase separation in vitro and in cells ^13,17,19–21^. These proteins interact with other nucleolar components, such as rRNA, snoRNAs, and/or ribosomal proteins, to form liquid-like condensates ^22^. The spatial heterogeneity observed within the nucleolus arises from the compartmentalization of different nucleolar proteins and their associated biomolecules into distinct condensed phases ^13^. The rRNA processing occurs in distinct spatial layers within the nucleolus, and specific defects in rRNA processing can disrupt the normal layered organization of nucleolar phases ^23^. Various factors, such as concentration, post-translational modifications, and cellular state, influence the phase separation properties of nucleolar proteins. Changes in these factors can modulate biomolecular interactions and phase separation, leading to the formation or dissolution of condensed phases within the nucleolus ^8^. However, an understanding of the internal structure and dynamics of the individual sub-compartments of the nucleolus, e.g., the GC, through the lens of phase separation remains incomplete.

Recent advances in super-resolution microscopy have created opportunities for investigating the sub-structural organization of nucleolar compartments ^24^. However, how the structural heterogeneity in the GC contributes to the multiple steps of ribosome subunit assembly ^15^ remains unexplained. For example, the GC has its name because it appears to comprise disorganized, heterogeneous granules in electron microscopy images ^25,26^. These granules are thought to contain rRNA, ribosomal proteins, and protein factors involved in the ribosome subunit assembly ^12,27,28^. Given the roles of differential biomolecular composition and material properties in the hierarchical organization of the primary nucleolar sub-compartments, we hypothesized that the spatial heterogeneity of the GC arises from sub-phases comprised of distinct subsets of its biomolecular components. To test this hypothesis, we considered the behavior of three key GC components: rRNA, which forms the catalytic core of ribosomes; NPM1, the highly abundant multivalent scaffold protein ^19^; and surfeit locus protein 6 (SURF6), an essential ribosomal subunit assembly factor ^15^. NPM1 establishes the liquid-like environment of the GC through phase separation, while SURF6 is an arginine (Arg)- and lysine (Lys)-rich nucleolar protein that prevents premature folding of key rRNA elements during pre-60S biogenesis ^15^. NPM1 was previously shown to undergo phase separation individually with rRNA and SURF6 ^17,19,21^, but the behavior of more complex mixtures was unknown, although a truncated form of SURF6 termed SURF6-N was previously shown to form homogeneous ternary condensates with NPM1 and rRNA ^18^. Here, we first probed the behavior of these three components in the setting of the GC using structured illumination microscopy (SIM), which revealed preferential colocalization of rRNA with SURF6 and partial spatial delocalization of rRNA from NPM1. While in vitro model systems have revealed the general role of phase separation in the formation of the GC ^17–19^, the origins of spatial heterogeneity and its contribution to ribosome subunit assembly remain elusive. Here, we developed a novel in vitro model system that recapitulates the GC sub-phases observed in cells. In our in vitro system, NPM1, SURF6, and rRNA form condensates with two coexisting dense phases that reflect the spatial heterogeneity observed in the GC. These so-called multiphase condensates exhibit core-shell architecture, with rRNA and SURF6 enriched in the core phase, which is hydrophobic, and NPM1 enriched in the shell phase, which is hydrophilic. The core and shell phases retain liquid-like material properties and dynamically rearrange when conditions are changed. Varying component concentrations dynamically modulated multiphase behavior both in vitro and in nucleoli. Importantly, we observed that the packaging of rRNA with ribosomal proteins weakens interactions with SURF6, enabling NPM1 to extract assembled ribosomes from the core condensate phase. Our results indicate that, as it assembles with ribosomal proteins, rRNA flows between at least two GC sub-phases enriched in SURF6 and NPM1, respectively, en route to the nucleoplasm. Sequence analyses indicate that many additional assembly factors can interact with rRNA similarly to SURF6, creating a GC sub-phase for early ribosomal subunit assembly steps. These findings reveal how spatial heterogeneity, arising from a complex interplay of molecular interactions within the GC, facilitates ribosome subunit assembly.

## Results

### Super-resolution imaging reveals the molecular underpinnings of spatial heterogeneity in the nucleolar GC

While long known to be structurally heterogeneous, comprising ribonucleoprotein (RNP) complexes ^9,10,25,26^, the physical origins of the granularity of the GC of the nucleolus have remained elusive. The length scale of the granular structure of the GC is below the traditional light microscopy diffraction limit. Therefore, we used SIM, a super-resolution fluorescence imaging technique ^29,30^, to resolve nucleoli of human DLD-1 colorectal adenocarcinoma cells engineered using CRISPR-Cas9 to express endogenous NPM1 tagged with monomeric, enhanced green fluorescent protein (mEGFP) and FBL tagged with a monomeric red fluorescent protein (mCherry; DLD-1^NPM1-G/FBL-R^ cells) ^31^. We used these cells to probe NPM1 as a GC maker, FIB for the DFC, and upstream binding factor (UBF) for the FC (Figures 1A and S1A; see Method Details), which revealed the classical hierarchical nucleolar structure, with the structural heterogeneity of NPM1 within the GC evident.

**Figure 1.**
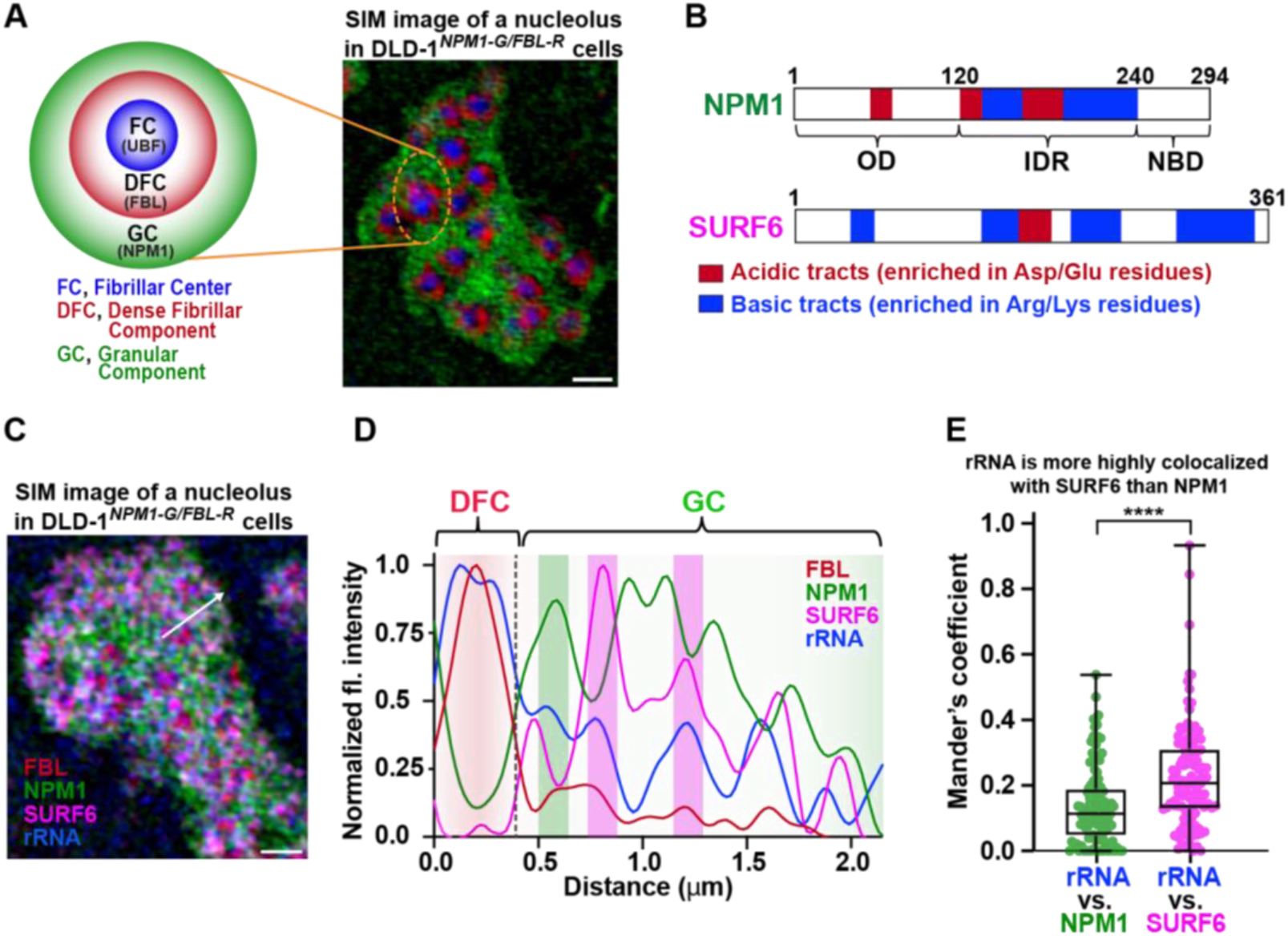
Super-resolution imaging resolves the molecular basis for spatial heterogeneity in the GC. (A) Schematic illustration of the three-layered structure of a nucleolus (left). The innermost central layer is the fibrillar center (FC), surrounded by the dense fibrillar component (DFC) embedded in the outermost granular component (GC). 2-D Structured illumination microscopy (SIM) image of DLD-1^NPM1-G/FBL-R^ cells immunostained to probe NPM1 (green) and FBL (red) with anti-GFP and anti-RFP nanobodies conjugated to AF488 and AF568, respectively. UBF (blue, pseudo color) was probed as an FC marker using an anti-UBF-AF647 antibody, scale bar = 1 μm. (B) Domain architecture of NPM1 (top) and SURF6 (bottom). NPM1 contains an oligomerization domain (OD), an intrinsically disordered region (IDR), and a nucleotide-binding domain (NBD). The OD contains an acidic tract (red), and the IDR is comprised of two acidic tracts (red) interspersed with basic tracts (blue). SURF6 is composed of a predominantly disordered N-terminal and a helical region in the C-terminal and exhibits four Arg-rich basic tracts (blue) and a single acidic tract (red). (C) SIM image of a nucleolus of a DLD-1^NPM1-G/FBL-R^ cell immunostained for SURF6 (magenta) and probed for newly synthesized rRNA with 5-ethynyl uridine (EU)/AF647 labeling (blue), in addition to immunostained for NPM1-mEGFP (green) and mCherry-FBL (red), scale bar = 1 μm. See Method Details. (D) Representative normalized fluorescence intensity line plot across the image in panel C (white arrow) for nucleolar components, FBL (red), NPM1 (green), SURF6 (magenta), and rRNA (blue). The green bar highlights a region enriched in NPM1 and rRNA, and the magenta bars highlight regions enriched in SURF6 and rRNA. (E) Plot of the degree of colocalization of rRNA versus NPM1 and rRNA versus SURF6 calculated from segmented DFC with surrounding inner GC regions. Mander’s overlap coefficient values are shown for 112 DFC/GC regions analyzed in six nucleoli; the two-tailed paired t-test was applied, (****, p < 0.0001).

To gain insight into the molecular basis of heterogeneity in the GC, we used SIM to image FBL in the DFC and NPM1, SURF6, and rRNA in the GC. NPM1 is an abundant, multivalent GC scaffold protein that undergoes phase separation with ribosomal and non-ribosomal proteins, including SURF6 and mediates molecular hand-offs that facilitate ribosome subunit assembly and efflux from the nucleolus ^17–19,21^. NPM1 displays a pentameric N-terminal oligomerization domain (OD), a central intrinsically disordered region (IDR) with alternating acidic and basic tracts, and a C-terminal nucleic acid binding domain (NBD) (Figures 1B, top and S1B). Acidic tracts in the NPM1 IDR mediate binding to Arg-rich basic tracts in other proteins, including SURF6, and the NBD mediates binding to rRNA (Figure S1B) ^17,19,21^. SURF6 is an essential Arg- and Lys-rich disordered protein displaying multiple basic tracts (Figures 1B, bottom and S1B) that functions as an early assembly factor in the maturation of pre-60S ribosomal subunits ^15^. While FBL, NPM1 and SURF6 were visualized using immunofluorescence (RB2AF568 antibody for FBL, GB2AF488 antibody for NPM1, and anti-SURF6-Dylight405 antibody for SURF6; see Method Details), newly transcribed rRNA was visualized using metabolic labeling with 5-ethynyluridine (EU) and Click chemistry with an Alexa-647 fluorophore ^32^. SIM images of these four biomolecules further revealed the structural heterogeneity of the nucleolus (Figures 1C and S1C), with the fluorescence intensity profile for a vector extending from one DFC through the GC to the nucleolar periphery revealing heterogeneous localization of NPM1, SURF6, and rRNA in the GC (Figure 1D). This structural heterogeneity was not resolved in our prior studies of nucleolar NPM1 and SURF6 due to the use of conventional confocal fluorescence microscopy ^18^. Because SURF6 is an early assembly factor in pre-60S biogenesis, which begins with rRNA species that emerge from the DFC into the GC, we performed this analysis within segmented DFC and immediately surrounding GC regions (Figure S1D). We quantified the apparent anti-correlated localization of rRNA versus NPM1 and SURF6 by computing the Mander’s overlap coefficient for rRNA and NPM1 fluorescence and, separately, rRNA and SURF6 fluorescence, in regions including and surrounding multiple DFCs within multiple nucleoli, which revealed that SURF6 is significantly more highly colocalized with rRNA than is NPM1 (Figures 1E, S1E, and S1F). These results indicate that differential localization of NPM1, SURF6, and rRNA underlie the structural heterogeneity of the GC of the nucleolus. Next, we investigated the mechanistic origins of this heterogeneity using in vitro reconstitution with the three GC biomolecules.

### Competing interactions between NPM 1, SURF6, and rRNA create spatial heterogeneity within multiphase condensates

NPM1, SURF6, and rRNA coexist within the GC through phase separation mediated by different multivalent interactions. We previously showed that acidic tracts in the intrinsically disordered region (IDR) of NPM1 interact with the Arg-rich motifs in other proteins, including SURF6 (Figure 1B), and the nucleic acid binding domain interacts with the rRNA ^17,19,21^. We also showed that the Arg- and Lys-rich N-terminal IDR of SURF6 interacts with rRNA and the acidic tracts of NPM1 ^17^. To quantify pair-wise interactions of these components, we generated binary phase diagrams using confocal fluorescence microscopy (Figures 2A-2C and S2A-S2C). NPM1 and SURF6 were visualized through conjugation with Alexa Fluor 488 (NPM1-AF488) and Alexa Fluor 647 (SURF6-AF647), respectively, and rRNA was visualized using SYTO 40 dye (Figure S3). SURF6 exhibited homotypic phase separation and heterotypic phase separation with NPM1 and rRNA and formed heterotypic condensates with rRNA at a 10-fold lower concentration than did NPM1 (Figures 2A-2D and S3). Next, we chose the concentrations marked by yellow circles in the phase diagrams (Figure 2A-2C) to test the behavior of a ternary mixture of the components. Strikingly, we observed the formation of multiphase condensates that exhibit core-shell architecture (Figure 2E). Analysis of fluorescence intensities using the Pearson correlation coefficient (PCC) revealed that rRNA localization is correlated with that of SURF6 and anti-correlated with that of NPM1 (Figure 2F), with rRNA and SURF6 6-fold and ⁓3-fold enriched in the core, respectively, and NPM1 ⁓4-fold enriched in the shell (Figures 2G and 2H). We next performed time-lapse imaging and observed that shells fuse within seconds while cores coalesce on the timescale of minutes (Figure 2I). The fused shells and cores transitioned from initial dumbbell-like morphology to a relaxed near-spherical shape over time (Figure 2I). Rapid fusion of shell phases preceded slower fusion of cores (Video S1). Together, our results show that SURF6 mediates the formation of liquid-like ternary multiphase condensates through phase separation involving homotypic interactions and differential heterotypic interactions with rRNA and NPM1. To understand the driving forces underlying their core-shell architecture, we next investigated the physicochemical properties of the two phases of reconstituted multiphase condensates.

**Figure 2.**
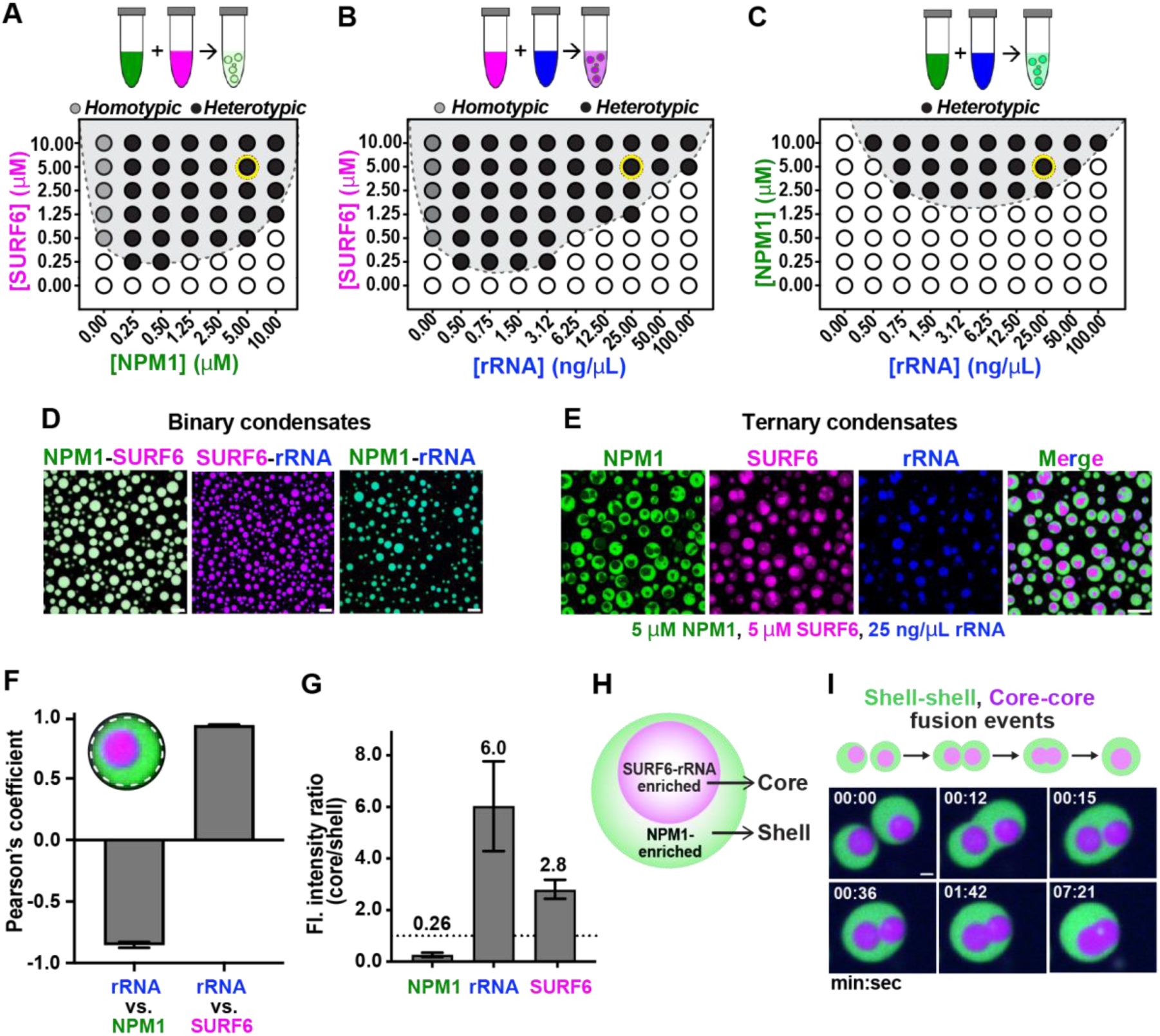
Competing interactions between NPM1, SURF6, and rRNA drive formation of multiphase condensates. LLPS phase diagrams for (A) NPM1-SURF6, (B) SURF6-rRNA, and (C) NPM1-rRNA mixtures in 10 mM Tris, 150 mM NaCl, 2 mM DTT, pH 7.5 buffer. The shaded gray regions indicate conditions with observed condensates using fluorescence confocal microscopy (see Figure S2 for images). Light and dark gray circles indicate homotypic and heterotypic phase separation, respectively. Phase separation was not observed under conditions marked by open circles. (D) Representative images of merged channels for binary condensates formed for NPM1-SURF6, SURF6-rRNA, and NPM1-rRNA at concentrations indicated by the yellow circles in panels A-C, respectively, scale bar = 10 µm. NPM1 was conjugated with AF488 (green) and SURF6 with AF647 (magenta). rRNA was stained with SYTO 40 (blue). (E) Fluorescence micrographs of ternary multiphase condensates of NPM1-SURF6-rRNA formed at concentrations used in panel D, scale bar = 10 µm. The multiphase condensates exhibited core-shell architecture. (F) Pearson’s correlation coefficient values for rRNA vs. NPM1 and rRNA vs. SURF6 calculated from the entire multiphase condensates, as illustrated in the inset. Values represent mean ± SD of 248 cores and 254 shell regions. (G) Plot of NPM1, rRNA, and SURF6 fluorescence intensity ratios inside the core and shell. Values represent mean ± SD of 370 condensates. (H) Illustration of a multiphase condensate indicating enrichment of SURF6-rRNA in the core and NPM1 in the shell. (I) Time-lapse microscopy images of representative fusion events of shells and cores, scale bar = 1 µm. Shells fuse more rapidly than cores. Also, see Video S1.

### Disparate physicochemical properties underlie the core-shell architecture of multiphase condensates

Nucleolar sub-compartments have different material and physiochemical properties due to their different biomolecular compositions ^13^. These properties, including surface tension, diffusivity, and viscoelasticity ^13,33^, influence the dynamic exchange of components within the nucleolus. In multiphase condensates, with SURF6-rRNA forming the core, we hypothesized that intrinsic properties of the core phase and its components underlie the hierarchical core-shell architecture. To test this hypothesis, we separately formed mimics of the core (SURF6-rRNA) and shell (NPM1-SURF6) and combined them (Figure 3A). Interestingly, within minutes, the cores were engulfed by the shell phase (Figure 3B and Video S2). Next, we individually monitored the accumulation of added SURF6 or rRNA in the cores within pre-made multiphase condensates. To examine SURF6 accumulation, we initially formed condensates with unlabeled SURF6 and then, after 2 hours, added a small amount of fluorescently labeled SURF6 (fl-SURF6, 2% of the total SURF6; Figure 3C). The labeled SURF6 diffused through the shell phase, maintaining a low concentration, and steadily accumulated in the core phase, with the fluorescence intensity approximately 3-fold greater in the core versus the shell phase (Figures 3D, 3E, and Video S3). Likewise, to monitor rRNA accumulation, we added a three-fold excess of rRNA to condensates initially formed with a low concentration of rRNA (Figure 3F). Similarly, rRNA diffused through the shell phase and accumulated in the core phase, causing the core volume fraction to increase by approximately two-fold within one hour (Figures 3G, 3H, and S4). Collectively, these results indicate that the components of the cores produce a phase with greater surface tension than the shells, causing cores to reside within the shell phases. Further, our results show that the multiphase condensates are under equilibrium control by dynamically responding to changes in component concentrations.

**Figure 3.**
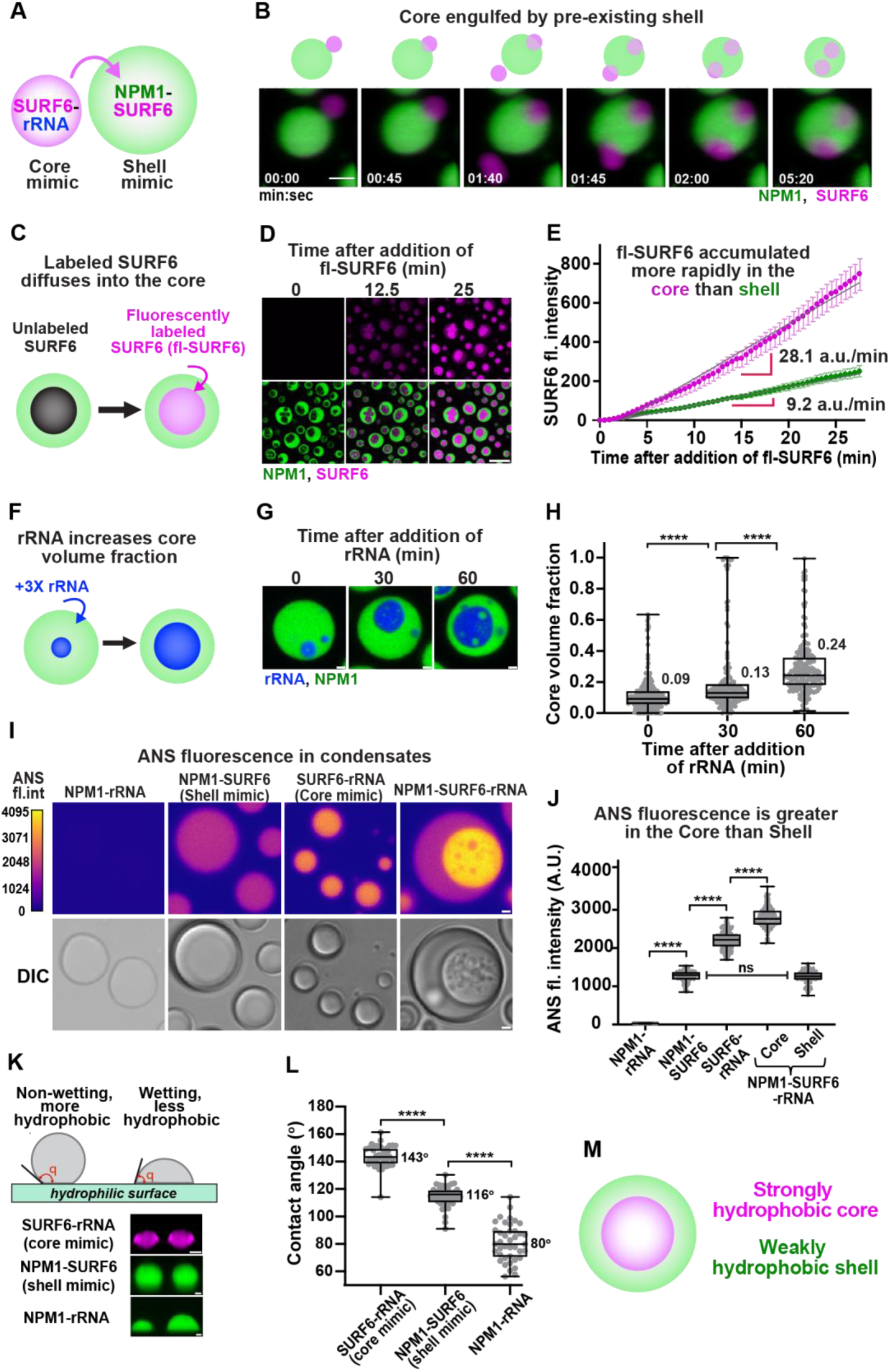
Emergence of core-shell architecture due to distinct physical properties of the nucleolar components. (A) Illustration of SURF6-rRNA condensates (core mimic) addition into pre-formed NPM1-SURF6 condensates (shell mimic). (B) Time-lapse lattice light-sheet microscopy (LLSM) images showing engulfment of cores by pre-existing shells, scale bar = 2 µm, SURF6 channel (magenta), and NPM1 channel (green), see Video S2. (C) Schematic representation of fluorescently-labeled SURF6-AF647 (fl-SURF6) accumulation into the cores of pre-formed multiphase condensates containing NPM1-AF488 and unlabeled SURF6. (D) Time-lapse images of fl-SURF6 accumulation into the cores of multiphase condensates. Top row (SURF6 channel) and bottom row (overlay of the NPM1 and SURF6 channels), scale bar = 10 µm, see Video S3. (E) Plot of fluorescence intensity of fl-SURF6 as a function of time, showing the rate of SURF6 accumulation in cores and shells. Values represent mean ± SD, n=12 condensates. (F) Schematic representation showing an increase in the volume fraction of cores upon the addition of excess rRNA (blue) into multiphase condensates. (G) Images of NPM1 and rRNA in multiphase condensates forming larger core phases as rRNA concentration increases, scale bar = 10 µm. (H) Plot of the increase in volume fraction of cores over time after adding 3-fold excess of rRNA, n = 243, 160 and 184 condensates for 0 min, 30 min and 60 min time-points. (I) Fluorescence images of condensates stained with 8-Anilino-1-naphthalenesulfonic acid (ANS) dye pseudo-colored with mpl-plasma LUT (top) and DIC (bottom) for binary (NPM1-rRNA, NPM1-SURF6, SURF6-rRNA) and ternary (NPM1-SURF6-rRNA) condensates, scale bar = 10 µm. (J) Quantification of ANS fluorescence intensity for binary (n = 115, 149, 208) and ternary (n = 95, 115) condensates, two-tailed t-test was applied, (****, p < 0.0001). (K) Schematic for determination of contact angle measurements (top). XZ projections of LLSM images for binary condensates on coverslips with a hydrophilic coating (bottom), scale bar = 2 µm. (L) Contact angles measured for binary condensates on a hydrophilic surface, n = 40 condensates, two-tailed Mann-Whitney test was applied (****, p < 0.0001). (M) Illustration of distinct physical properties of the core and shell phases of multiphase condensates.

We next investigated the origins of the apparent core-shell surface tension differences, which we hypothesized are due to differences in the hydrophobicity of the two phases, as we showed previously for the DFC and GC ^13^ and preliminarily showed for NPM1/SURF6-N condensates ^19^. To test this idea, we used 8-anilinonaphthalene-1-sulfonic acid (ANS), a fluorescent dye that binds to hydrophobic pockets in partially folded molten globule proteins ^34,35^, to probe the physicochemical character of the core and shell phases. ANS is also known to bind Arg and Lys residues in proteins through ion pair interactions ^36,37^. Fluorescence microscopy and bright-field microscopy (Figure 3I, top and bottom, respectively) of various combinations of unlabeled NPM1, SURF6, and rRNA revealed significantly enhanced ANS fluorescence in the cores compared to shells of ternary multiphase condensates and in binary core versus shell mimetic condensates. ANS fluorescence intensity was greatest in multiphase condensate cores and binary cores mimetic condensates, which are enriched in SURF6 and rRNA, and significantly lower in the corresponding shell phases, which are enriched in NPM1 (Figures 2G and 3J). We next investigated the surface properties of these condensates using lattice light-sheet microscopy to measure their contact angles on a hydrophilic surface (Figure 3K), with the expectation that hydrophobic condensates would resist wetting. Consistent with the results using ANS dye, the mean contact angle was greatest for core mimetic condensates (143⁰), with shell mimetic condensates exhibiting smaller contact angles (116⁰) and NPM1-rRNA condensates displaying the smallest contact angles (80⁰) (Figure 3L). Together, the ANS dye and surface wetting results indicate that the core phases possess a hydrophobic character (Figure 3M) and have relatively high surface tension and, thus, are energetically favored to reside within the interior of multiphase condensates.

### Dynamic interplay between NPM 1, SURF6, and rRNA in vitro and in cells

Our previous “hand-off” model proposed that NPM1 facilitates ribosome biogenesis by independently undergoing phase separation with rRNA and ribosomal proteins, which enter the GC via opposing fluxes, creating a liquid-like environment for ribosomal subunit assembly ^17,18^. Here, we revisited this model based on our new findings, showing that the GC protein SURF6 preferentially binds rRNA in the presence of NPM1. We first titrated SURF6 into mixtures with fixed concentrations of NPM1 and rRNA (5 μM and 25 ng/μL, respectively). At SURF6 concentrations up to 2.5 μM, homogenous ternary condensates were observed. However, above this concentration, the condensates decomposed into two dense phases with the core - shell architecture noted above (Figures 4A, upper row and S5A). Up to 2.5 μM, SURF6 progressively displaced NPM1 from the homogeneous condensates into the surrounding light phase, decreasing the apparent NPM1 partition coefficient (K_app_; Figures 4A, lower row, 4B, and S5A). Above 2.5 μM, most SURF6 formed the core phase with rRNA, while a small portion outside the cores underwent phase separation with NPM1 to form the shell phase. As the SURF6 concentration increased, NPM1 became increasingly delocalized from rRNA (Figure 4C). These results show that SURF6 preferentially interacts with rRNA, forming a hydrophobic phase that is incompatible with the surrounding rRNA-depleted, hydrophilic NPM1-rich shell phase (Figure 4D).

**Figure 4.**
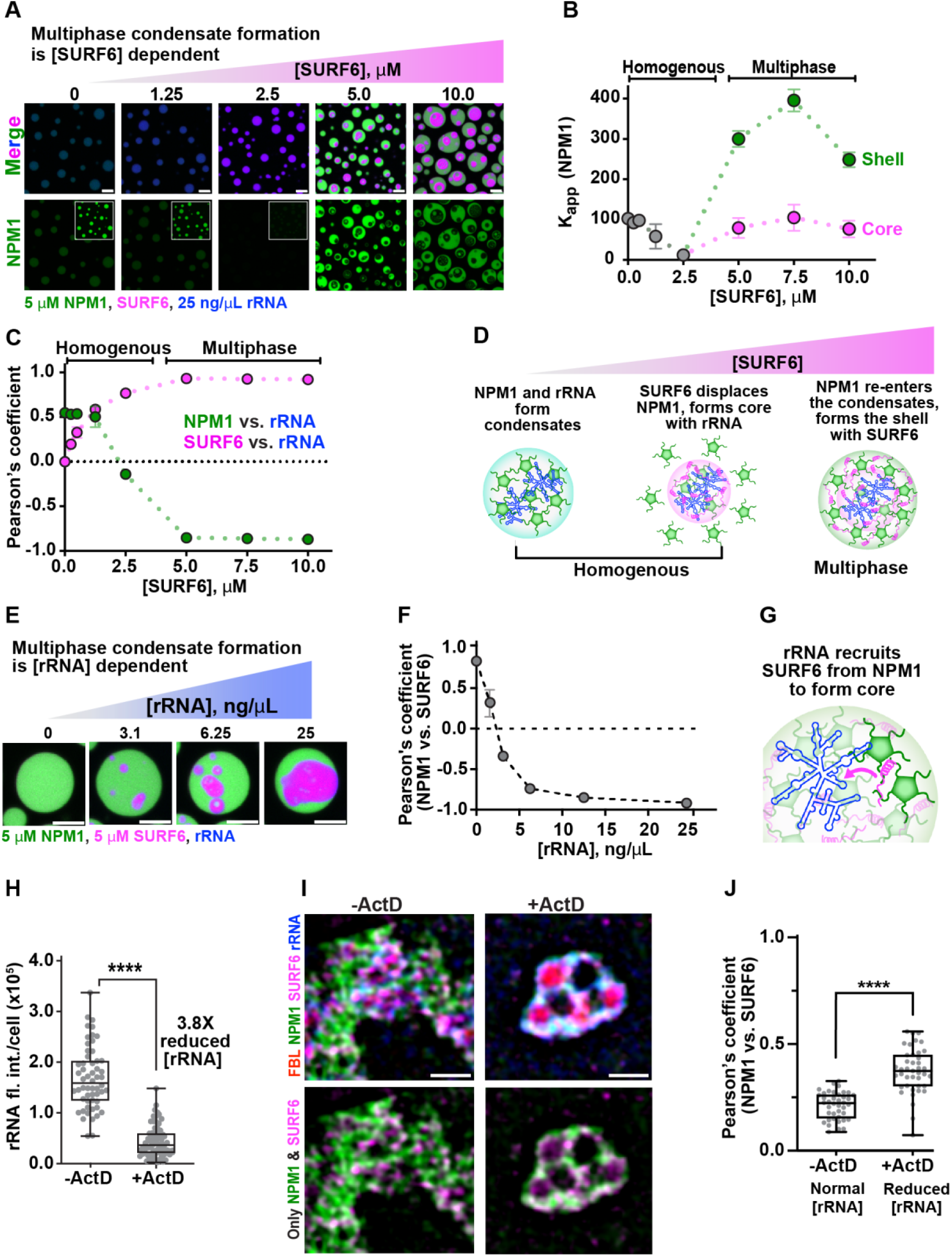
Concentration-dependent effect of SURF6 and rRNA on the multiphase condensate formation and spatial heterogeneity within GC. (A) Confocal fluorescence microscopy images of condensates formed upon titration of SURF6 (magenta) into pre-formed NPM1 (green) and rRNA (blue) condensates, scale bar = 5 µm. Pre-incubated condensates containing 5 μM NPM1 and 25 ng/μL rRNA were titrated with SURF6 and imaged after three hours. The fluorescence intensity threshold for the inset images in the bottom row was reduced 4-fold. (B) Partition coefficient (K_app_) values for NPM1 as a function of increasing SURF6 concentration. (C) Pearson’s correlation coefficient for NPM1 vs. rRNA and SURF6 vs. rRNA as a function of SURF6 concentration. Values for panels B and C represent mean ± SD, n ≥ 107 condensates. (D) Schematic illustration of the molecular mechanism of formation of multiphase condensates with increasing SURF6 concentration. (E) Confocal fluorescence microscopy images of condensates formed upon titration of additional rRNA into pre-formed condensates containing 5 μM NPM1 and 5 μM SURF6, scale bar = 5 µm. (F) Pearson’s correlation coefficient values for NPM1 vs. SURF6 as a function of increasing rRNA concentration. Values represent mean ± SD, n ≥ 187 condensates. (G) Schematic diagram illustrating the formation of multiphase condensates with increasing rRNA concentration leading to larger SURF6-rRNA cores as rRNA recruits more SURF6 into cores. (H) Sum of nucleolar rRNA fluorescence intensities quantified per cell in actinomycin D (ActD) treated (n=70) and untreated (n=56) DLD-1^NPM1-G/FBL-R^ cells, two-tailed Mann-Whitney test was applied, (****, p < 0.0001). (I) Representative SIM images of untreated and ActD treated DLD-1^NPM1-G/FBL-R^ cells, scale bar = 10 µm. The top row of images are overlaid channels of FBL (red), NPM1 (green), SURF6 (magenta), and rRNA (blue) in the absence (top, left) and presence (top, right) of Act D. The bottom row shows overlaid images for only the NPM1 (green) and SURF6 (magenta) channels. (J) Pearson’s correlation coefficient values for NPM1 versus SURF6 for ActD-treated (n = 41) and untreated cells (n = 40), two-tailed Mann-Whitney test was applied, (****, p < 0.0001).

To gain insight into the effect of rRNA concentration on multiphase condensate formation, we next titrated rRNA into homogeneous condensates formed with equal concentrations of NPM1 and SURF6 (5 μM each). Upon first addition, rRNA was sequestered by SURF6 within core phases, which increased in size as the rRNA concentration was increased (Figures 4E and S5B). As seen with the titration of SURF6, NPM1 became increasingly delocalized from the SURF6-rich core phase within multiphase condensates as the rRNA concentration increased (Figures 4F and 4G). These results demonstrate that SURF6 has a high capacity to sequester nascent rRNA within a dense, liquid-like hydrophobic phase in the presence of equimolar NPM1.

Our in vitro results showed that interactions between SURF6 and NPM1 within condensates are regulated by rRNA, with the two proteins co-localized at low rRNA concentrations. To explore this regulatory interplay in cells, we treated DLD-1^NPM1-G/FBL-R^ cells with Actinomycin D (ActD) to halt RNA polymerase I-dependent transcription ^38–40^ and reduce the level of rRNA (Figures 4H and S5C). While treatment with ActD is well appreciated to induce the formation of so-called nucleolar caps comprised of FC and DFC components ^41^, the effects of this treatment on the remaining GC components were unknown. NPM1 and SURF6 were partially spatially delocalized within the GC of DLD-1^NPM1-G/FBL-R^ cells in the absence of ActD treatment (Figures 4I (left), 4J, S5D, and S5E). However, when rRNA was reduced through ActD treatment, spatial colocalization of NPM1 and SURF6 increased as quantified using the PCC (Figures 4I (right), 4J, S5D, and S5E), as was observed in vitro, with the PCC value for NPM1 and SURF6 colocalization reaching higher values at lower rRNA concentrations. Together, our in vitro and cellular results demonstrate the dynamic interplay between NPM1, SURF6, and rRNA in the nucleolus mediated by multiphase behavior that gives rise to the GC’s namesake granularity. Next, we investigated the implications of this dynamic interplay in the flux of assembled ribosomal subunits out of the GC.

### The dynamic interplay between nucleolar components in the GC regulates the outward flux of ribosomal subunits

We previously showed that the flux of rRNA within assembled ribosomal subunits out of the GC is facilitated by reduced thermodynamic favorability of interactions with NPM1 that underlie phase separation ^18^. As rRNA assembles with ribosomal proteins within subunits, interaction sites are buried, weakening interactions with NPM1. However, at the time of this report, the role of SURF6/rRNA interactions in creating the heterogeneous structure of the GC and their implications for the outward flux of assembled ribosomal subunits were not appreciated. Therefore, we revisited this “flux model” through the lens of multiphase NPM1/SURF6/rRNA condensates. We first showed that SURF6 has a greater propensity to phase separate with unassembled, nascent rRNA than rRNA within assembled ribosomes, forming more, larger, and denser condensates with the former rRNA species (Figures 5A-5C, S6A, and S6B). We next separately formed multiphase condensates with NPM1, SURF6, and these two rRNA species and then added a stoichiometric excess of NPM1 to mimic the environment of the GC, in which NPM1 is highly abundant ^17,18^. Most unassembled rRNA (77%) was retained within condensates after addition of excess NPM1, although the change in conditions eliminated the core-shell architecture (Figures 5D (left), 5E (left), and S6C). In contrast, only 28% of rRNA assembled within ribosomes was retained in condensates (Figures 5D (right), 5E (right), and S6D), indicating that NPM1 effectively extracted assembled rRNA from the SURF6-rich core of multiphase condensates. Using the K_app_ values for the two pairs of conditions, we determined that extraction of assembled rRNA from multiphase condensates by NPM1 is significantly more thermodynamically favorable (free energy of rRNA exit, ΔΔG^Ef f lux^ = -0.43 kcal/mol; see Method Details) than for unassembled rRNA (ΔΔG^Ef flux^ = -0.12 kcal/mol; Figures 5F, S6E, and S6F). These findings highlight the role of interplay between SURF6 and NPM1 and their interactions with rRNA in mediating the retention of nascent, unassembled rRNA within GC-like multiphase condensates and gating the exit of mature, assembled rRNA from them (Figure 5G).

**Figure 5.**
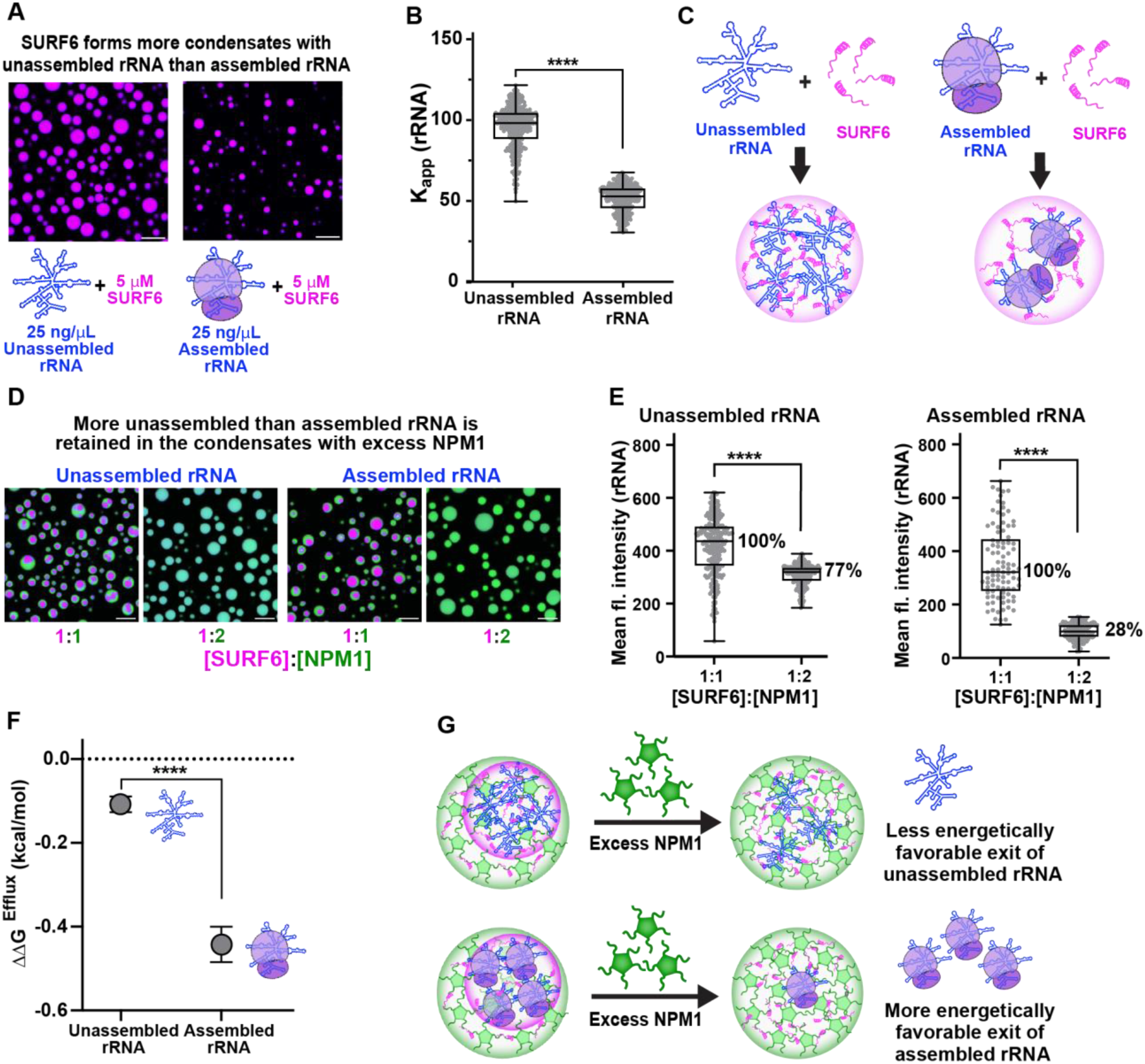
Dynamic interplay between nucleolar components in the GC regulates the outward flux of ribosomal subunits. (A) Fluorescence micrograph of condensates comprised of 5 μM SURF6 with 25 ng/μL unassembled rRNA (total rRNA, left) or 5 μM SURF6 with 25 ng/μL assembled rRNA (70S ribosomes, right) in 10 mM Tris, 150 mM NaCl, 2 mM DTT, 3 mM MgCl2, pH 7.5 buffer. Total rRNA and 70S ribosomes are labeled with SYTO 40 dye, scale bar = 10 µm. (B) Comparison of the partition coefficient (K_app_) values for rRNA (stained with SYTO 40) in the condensates formed in panel A, n = 353 and 446 condensates for unassembled rRNA and assembled rRNA, respectively. (C) Schematic depicting reduced partitioning of assembled rRNA into SURF6-rRNA condensates compared to unassembled rRNA. (D) Fluorescence micrographs of condensates comprised of NPM1 (green), SURF6 (magenta), and unassembled (left) or assembled (right) rRNA (blue) at 1:1 and 1:2 concentration ratios of SURF6:NPM1, scale bar = 10 µm. (E) Plot of SYTO 40 mean fluorescence intensity of condensates at the two different concentration ratios of SURF6:NPM1 containing unassembled rRNA (left, n = 229, 224) and assembled rRNA (right, n =93, 143), two-tailed t-test was applied, (****, p < 0.0001). (F) Calculated Gibbs free energies of particle exit (ΔΔG^Efflux^) of unassembled and assembled rRNA from the multiphase condensates from panel E upon the addition of excess NPM1. (G) Schematic depicting a more favorable exit of assembled rRNA from multiphase condensates due to the addition of excess NPM1 as a consequence of weakened interactions with SURF6.

## Discussion

Our data show that structural heterogeneity in the GC arises through the formation of liquid-like sub-phases marked by the differential localization of NPM1, SURF6, and rRNA (Figure 6). We used super-resolution SIM, which has previously been used to study the structural organization of the nucleolus ^42,43^, to resolve the sub-compartmentalization and structural organization of biomolecules within the GC. Based on our findings, we propose that the early pre-60S assembly factor, SURF6, preferentially phase separates with nascent rRNA, confining it within an interior sub-phase of the dynamic GC. SURF6 is a non-ribosomal nucleolar protein that displays positively charged basic tracts that interact with negatively charged rRNA through electrostatic interactions. As ribosomal subunits assemble, rRNA migrates from the inner GC to the nucleolar periphery, while ribosomal proteins enter from the nucleoplasm into the GC ^17^. During the assembly process, NPM1, the most abundant GC protein, participates in heterotypic interactions, including with rRNA, SURF6, and ribosomal proteins, which promote phase separation (Figure 6). We propose that NPM1, concentrated in the central GC, dynamically regulates the interplay of these nucleolar constituents through competitive interactions, causing the formation of multiple sub-phases. An extension of this is that the NPM1-rich liquid-like milieu can buffer the concentrations of nucleolar proteins and enable the orderly assembly of pre-ribosomal particles within the GC and their efflux from the nucleolus.

**Figure 6.**
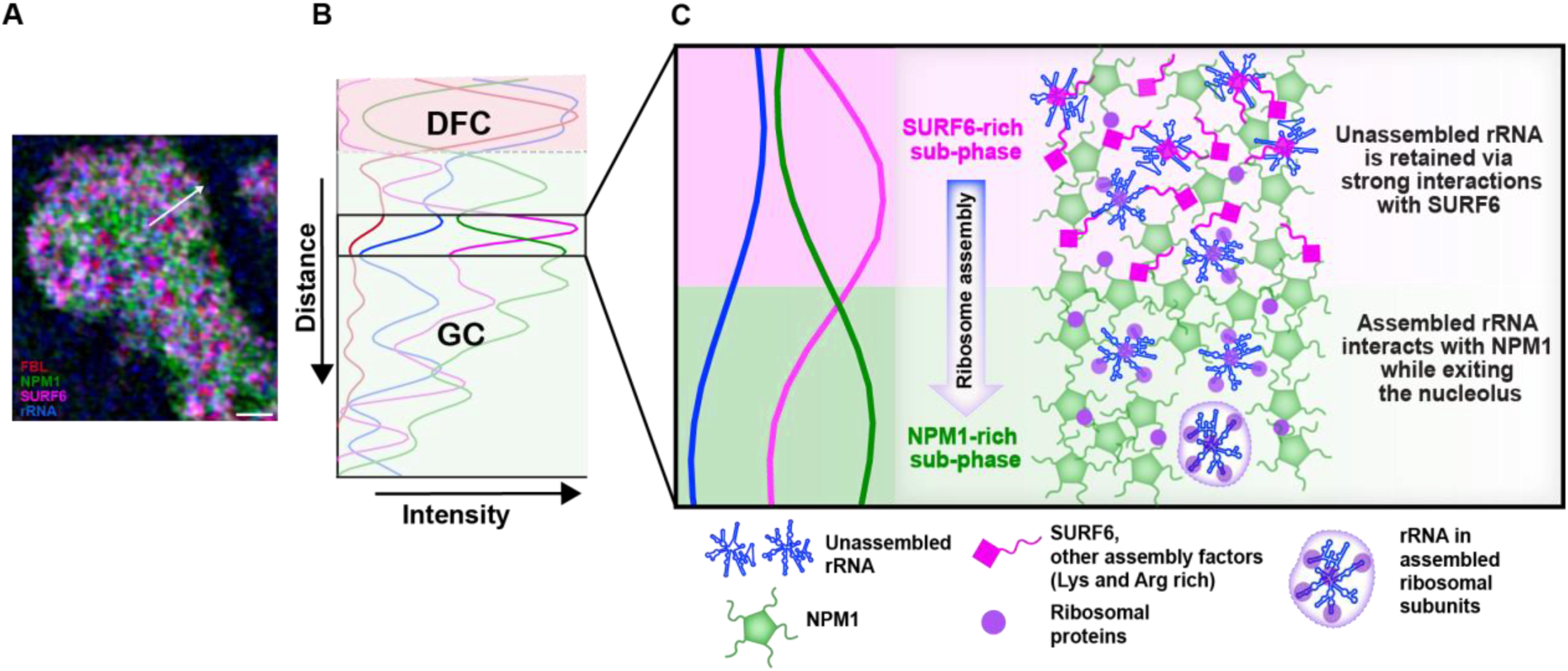
Structural heterogeneity in the GC of the nucleolus underlies ribosomal subunit assembly. Schematic diagram illustrating the organization of compositionally distinct, liquid-like SURF6-rich and NPM1-rich sub-phases within the nucleolar GC. (A) SIM image of a nucleolus (FBL, red; NPM1, green; SURF6, magenta; rRNA, blue). (B) A plot of normalized fluorescence intensities along the white arrow is shown in panel A, with the DFC and GC regions indicated. (C) The boxed region of the GC in panel B is expanded (left) to illustrate the spatial delocalization of SURF6 (magenta) and rRNA (blue) versus NPM1 (green). We propose (right) that, as ribosomal subunits assemble, they successively move outward (downward in the illustration) through SURF6-rich and NPM1-rich sub-phases under thermodynamic control mediated by differential interactions of these proteins with rRNA at different states of ribosomal subunit assembly. SURF6 prefers to bind rRNA in early assembly states, which enables highly abundant NPM1 to extract assembled ribosomal subunits from interior SURF6-rich sub-phases.

NPM1 is a multifunctional protein that, in addition to mediating ribosome biogenesis, plays roles in nuclear stress responses ^44^. We previously elucidated the mechanisms that govern the localization of NPM1 within nucleoli ^21^. These include interactions of acidic tracts within NPM1’s central IDR (Figure 1B, top) with tracts of basic residues, preferentially Arg residues, in other nucleolar proteins, including ribosomal proteins and SURF6 ^17,45^. In addition, the C-terminal NBD, flanked by a basic tract at the C-terminus of the IDR, interacts with nucleic acids, including rRNA. The valence of these two types of interacting regions is enhanced by the pentamerization of NPM1’s OD, promoting phase separation with oppositely charged substrates, basic proteins, and negatively charged rRNA. Here, we show that SURF6, comprised of 13% Lys and 15% Arg residues, competes with NPM1 for rRNA, initially ejecting it during titrations from homogeneous condensates comprised of NPM1 and rRNA and forming dense SURF6/rRNA-rich condensates. Upon further titration, however, due to its high capacity for multivalent interactions, NPM1 undergoes phase separation with the low concentration of SURF6 in the light phase that is in equilibrium with the SURF6/rRNA-rich condensates, forming the outer shell phase of multiphase condensates with SURF6/rRNA cores. Our results show that this architecture arises due to the hydrophobic character of the SURF6-rich cores and the hydrophilic character of the NPM1-rich shells. We ascribe the hydrophobic character of cores to the enrichment of SURF6 in Arg residues. While the Arg side chain displays dispersed positive charge, which interacts favorably with negatively charged nucleic acids such as rRNA, Arg residues in the poly-Arg peptide, protamine, were previously shown to mediate the formation of viscoelastic, hydrophobic condensates under conditions mimicking the crowded cellular environment ^46^. In contrast, basic tracts interspersed with the acidic tracts of NPM1’s IDR are enriched in Lys residues (24 Lys and 3 Arg residues), accounting for the hydrophilic character of the NPM1-rich shell phase in multiphase condensates. The high Arg content also mediates the high affinity of SURF6 for rRNA, enabling it to out-compete Lys-rich NPM1, as was discussed previously based on results using Arg- and Lys polymers ^47^. Additionally, its high Arg content may limit the solubility of SURF6 and contribute to the formation of the core phase within multiphase condensates. Thus, differences in the balance of Arg and Lys residues in SURF6 and the IDR of NPM1 mediate both the competitive preference of SURF6 for rRNA and the hydrophobic character of SURF6/rRNA sub-phases, which in turn govern the core-shell architecture of ternary, multiphase condensates with NPM1. We propose that these evolutionarily tuned amino acid biases underlie the preferential localization of SURF6 with rRNA in the GC region of the nucleolus.

SURF6, as it undergoes phase separation in the GC, also acts as an assembly factor in the early stages of pre-60S ribosome biogenesis ^15,48^. Recent cryogenic electron microscopy (cryo-EM) findings show that the C-terminal region of SURF6 (residues 212-361) adopts α-helical structure upon binding, together with SSF1 and RRP15, a specific region of rRNA within states A and B during the assembly of the pre-60S particle ^15,49^. The SURF6-SSF1-RRP15 complex organizes helices of the six structural domains of 28S rRNA, preventing premature folding of key rRNA structural elements. As assembly progresses to state C, the DEAD-box helicase DDX54 remodels the 28S rRNA, abolishing the binding site for the SURF6-SSF1-RRP15 complex, leading to its dissociation ^15^. These early assembly steps occur within the GC, with SURF6 and its assembly factor partners alternately associating with and dissociating from pre-60S particles as they assemble and mature. We propose that it’s the Arg and Lys residue-enrichment that enables SURF6 to be retained within the GC through phase separation as it cycles on and off pre-60S particles. Supporting this proposal, our data show that SURF6 undergoes phase separation in vitro with both protein-free rRNA and assembled ribosomes. Further, we previously showed that the intrinsically disordered N-terminal region of SURF6, SURF6-N (residues 1-182), undergoes phase separation with isolated NPM1 and rRNA ^18^. This region of SURF6, extending to residue 211 (Figure S7A), is unresolved in cryo-EM data for assembly states A and B, making it available for Arg and Lys residue-mediated nonspecific, multivalent interactions with immature rRNA species within the GC. Interestingly, two C-terminal regions of SSF1 (residues 205-265 and 297-473) are also structurally unresolved and likely disordered in the early assembly states, with the latter region enriched in Arg and Lys residues, similar to SURF6 (Figures S7A and S7B). Also, the N- and C-terminal regions of RRP15 are structurally unresolved (residues 1-126 and 212-282), and both are enriched in Asp and Glu residues interspersed with Arg and Lys residues (Figures S7A and S7B). The unresolved regions, disordered regions of SSF1 and RRP15 containing basic residues, may interact nonspecifically with rRNA similarly to the SURF6 IDR, while the IDRs with mixed acidic and basic character may interact with basic proteins in the GC, including assembly factors like SURF6 and possibly ribosomal proteins (Figure S7B). Many structurally characterized pre-60S assembly factors ^15^, in addition to SURF6, SSF1, and RRP15, have unresolved regions enriched in Arg and Lys residues (Figures S8A and S8B). Specifically, 28 of 36 assembly factors, from 12 structurally characterized assembly substates ^15^, display unresolved regions ≥ 30 residues in length that, on average, have Arg and Lys residue content similar to that of SURF6 and significantly greater that of the human IDRome, defined here as all IDRs in the human proteome ≥ 60 amino acids in length (Figures S8A and S8B). These results suggest that positively charged IDRs enable many pre-60S assembly factors to be retained in the liquid-like milieu of the GC through nonspecific interactions with 28S and other rRNA species. Large portions of these assembly factors adopt folded structures within specific pre-60S assembly states, which are ultimately disrupted and released as assembly progresses to yield mature 60S ribosomal subunits. Our observations highlight the synergy between constitutively disordered and conditionally folded regions of assembly factors that contribute to the liquid-like nature of the GC and facilitate the sequential assembly of specific structures within 60S ribosomal subunits.

As noted above, our SIM imaging data shows that the proteins, NPM1 and SURF6, differentially localize with rRNA in the GC, reflecting this nucleolar sub-compartment’s namesake granularity. This spatial heterogeneity arises from SURF6’s preference for binding to and phase separating with nascent rRNA, rooted in enrichment in Lys and, in particular, Arg residues within its largely disordered polypeptide chain. The dense islands of SURF6 and rRNA in the GC are surrounded by a sea of NPM1 phase separated with a low level of SURF6 and likely many other nucleolar proteins. Our in vitro data show that SURF6’s affinity for rRNA declines as ribosomes assemble and that, in the presence of excess NPM1, mature ribosomes can be extracted from multiphase condensate cores that they initially form in the presence of SURF6 and a reduced concentration of NPM1. NPM1 is among the most abundant proteins in human cells ^50^ and is largely concentrated through phase separation in the GC of nucleoli ^18,21^. SURF6, by contrast, is more than 50-fold less abundant in human cells ^50^ but, due to its physicochemical properties, is highly enriched through phase separation within nascent rRNA-rich sub-compartments in the GC. We propose that high affinity for nascent rRNA enables SURF6 to function as an assembly factor in the early stages of pre-60S subunit assembly and to be retained in the rRNA-rich sub-compartment within the GC. As pre-60S subunits progress through multiple assembly steps ^49^, SURF6’s affinity for them declines, enabling the subunits to migrate from SURF6-rich GC sub-phases into others containing highly abundant NPM1, which precedes their efflux from the nucleolus. In addition to undergoing phase separation with ribosomal components, NPM1 binds to assembled ribosomal subunits that have exited the nucleolus and shuttles with them to the cytoplasm, followed by recycling back to the nucleus ^51^. Thus, the interplay between SURF6, NPM1, ribosomal components, and numerous additional assembly factors, forming distinct, compositionally biased sub-phases, mediates the efflux of ribosomal subunits from the nucleolus coupled with their maturation. Broadly, these observations, made possible by super-resolution fluorescence imaging and in vitro modeling, illustrate how the formation of phase-separated mesoscale structures can orchestrate critical steps, e.g., ribosome subunit assembly, in the essential biological process of ribosome biogenesis.

## Method Details

### Cell lines

DLD-1 (male, adult, age not reported, Dukes’ type C colon cancer) cell lines were purchased from American Type Culture Collection (ATCC). DLD-1 cells were maintained in RPMI 1640 medium (Thermo Fisher Scientific), supplemented with 10% fetal bovine serum and 100 U/mL penicillin-streptomycin. Cultures were incubated at 37 °C in a humidified 5% CO₂ environment. Gene-edited lines underwent authentication through short tandem repeat (STR) profiling, and mycoplasma contamination was ruled out using the e-Myco PLUS Mycoplasma PCR Detection Kit (Bulldog Bio).

### Generation of endogenously NPM1 and FBL-tagged cell line

DLD1-NPM1-mEGFP/FBL-mCherry C-terminally double-tagged (DLD-1^NPM1-G/FBL-R^) cells were generated using CRISPR-Cas9 technology in the Center for Advanced Genome Engineering (St. Jude), as described^52^. Briefly, the DLD-1^NPM1-G/FBL-R^ cell line was created by transiently co-transfected 500,000 DLD1 cells with pre-complexed ribonuclear proteins (RNPs) consisting of 100 pmol of chemically modified sgRNA (CAGE117.NPM1.g1, Synthego), 33 pmol of 3X NLS *Sp*Cas9 protein (St. Jude Protein Production Core), and 500 ng of plasmid donor (Biobasic USA). The transfection was performed via nucleofection (Lonza, 4D-Nucleofector™ X-unit) using solution P3 and program CA-137 in a small (20 uL) cuvette according to the manufacturer’s recommended protocol. Single cells were sorted for GFP positivity five days post-nucleofection into 96-well plates containing prewarmed media and clonally expanded. Clones were screened and verified for the desired modification using PCR-based assays (5’ junction primers - CAGE117.gen.F2 and CAGE117.junc.meGFP.DS.R2, 3’ junction primers - CAGE117.junc.meGFP.DS.F2 and CAGE117.gen.R2, zygosity confirmation primers - CAGE117.DS.F and CAGE117.DS.R) and confirmed via sequencing. A resulting *NPM1 mEGFP* tagged clone (2C4) was then transiently co-transfected with CAGE223.FBL.g3, 3X NLS *Sp*Cas9 protein, and CAGE223.g3.mCherry.donor as described above. Additionally, 1 uM of Nedisertib (M3814) was added to the media for 24 hours after nucleofection. Cells were single-cell sorted for mEGFP and mCherry double-positive cells and clonally expanded. Clones were screened and verified for the desired targeted integration using PCR-based assays (5’ junction primers - CAGE223.gen.F and CAGE223.junc.DS.R, 3’ junction primers - CAGE223.junc.DS.F and CAGE223.gen.R, zygosity confirmation primers - CAGE223.FBL.DS.F and CAGE223.FBL.DS.R) followed by sequence confirmation. All final clones tested negative for mycoplasma by the MycoAlert^TM^Plus Mycoplasma Detection Kit (Lonza) and were authenticated using the PowerPlex® Fusion System (Promega) performed at the Hartwell Center (St. Jude). Editing construct sequences and screening primers are listed in Table S1.

### EU labeling and immunostaining

DLD-1^NPM1-G/FBL-R^ cells were grown to approximately 50% confluence on 12 mm diameter #1.5 coverslips and treated with 1 mM 5-ethynyl uridine (EU) for 30 minutes. Cells were fixed for 10 minutes using 4% paraformaldehyde (PFA) in PBS. Following fixation, coverslips were washed twice with Dulbecco’s phosphate-buffered saline (DPBS) buffer. Cells were permeabilized with 0.5% Triton X-100 in PBS for 5 mins. EU incorporated into newly synthesized RNA was fluorescently labeled with Alexa-647 fluorophore using an RNA Click-It RNA imaging kit (Molecular Probes) following the manufacturer’s protocol (https://assets.thermofisher.com/TFS-Assets/LSG/manuals/mp10329.pdf). For immunostaining, coverslips were initially blocked with 4% bovine serum albumin (BSA) in DPBS. To probe for SURF6, we used an anti-SURF6 rabbit polyclonal antibody (Invitrogen, PA5-54841) at 1:500 dilution. To obtain good-quality SIM imaging, we enhanced the signals of mEGFP in NPM1 and mCherry in fibrillarin using fluorophore-conjugated nanobodies. We accomplished this by adding GFP and RFP nanobodies conjugated to Alexa 488 and Alexa 568 fluorophores (GB2AF488, RB2AF568, Chromotek), respectively, to the SURF6 antibody mixture (1:500 dilution). Samples were incubated in the antibody mixture for an hour at room temperature. Bound SURF6 antibodies were probed using fluorophore-conjugated secondary antibody AffiniPure F(ab’)₂ Fragment Donkey Anti-Rabbit IgG1-Dylight 405 (Jackson ImmunoResearch, 711476152) diluted to 1:400. Coverslips were washed with DPBS and water and mounted on a glass slide using ProLong Diamond Anti-fade Mountant (Invitrogen). Samples were allowed to set for at least 72 hours prior to imaging.

To probe for the three layers of the nucleolus, we immunostained fixed DLD-1^NPM1-G/FBL-R^ cells with anti-GFP Alexa 488 (for NPM1) and anti-RFP Alexa 568 (for FBL) nanobodies (1:500 dilution) and anti-UBF conjugated to Alexa 647 (Santa Cruz Biotech, SC-13125) at 1:250 dilution. Mounted slides were similarly allowed to be set for 72 hours prior to imaging.

### Actinomycin D treatment

DLD-1^NPM1-G/FBL-R^ cells were seeded onto poly-Lysine coated coverslips to approximately 50% confluence and synchronized by overnight serum starvation. Media was replaced the following day, and cells were allowed to recover for two hours prior to 30 min treatment with 1 mM EU. Following this, cells were washed three times with pre-warmed media to remove residual EU. Post-wash, cells were incubated with media containing either 0.1 µg/mL actinomycin D (ActD) (Invitrogen) or DMSO for one hour. To minimize fixation artifacts, cells were directly fixed at the selected time point in media using 4% pre-warmed PFA.

### SIM imaging and processing

Multicolor super-resolution SIM (SR-SIM) images were collected using a Zeiss ELYRA PS.1 super-resolution microscope (Carl Zeiss MicroImaging) using a 100X Plan-Apochromat oil objective lens with 1.46 NA. Images were acquired using 405 nm (Dylight 405, SURF6), 488 nm (GFP/Alexa 488, NPM1), 561 nm (RFP/Alexa 568, FBL), and 660 nm (Alexa 647, rRNA) laser lines. Five orientation angles of the excitation grid were acquired for each Z plane, where Z spacing is 90 nm between planes. SIM images were processed using the SIM analysis module of the Zen 2012 BLACK software (Carl Zeiss MicroImaging). To image the three layers of the nucleolus, we used the 488 nm (GFP/Alexa 488, NPM1), 561 nm (RFP/Alexa 568, FBL), and 660 nm (Alexa 647, UBF) laser lines.

For proper spatial alignment of multichannel SR-SIM images, a post-acquisition channel alignment was performed for all SR-SIM images. To generate the calibration file, 100 nm TetraSpeck beads were seeded onto a #1.5 coverslip, mounted to a slide, and imaged using the same imaging parameters as the sample. Data was reconstructed with the SIM analysis module before using the Channel Alignment processing function, selecting the “Affine” and “Fit” options within the Zen 2012 BLACK software. After the calibration file was generated, it was saved and utilized for subsequent imaging experiments. SR-SIM images were aligned using the calibration file generated in the Zen 2012 BLACK software using the Channel Alignment processing function.

### Colocalization analyses

SIM images of nucleoli were visualized and analyzed using IMARIS software (Oxford Instruments). Due to the highly heterogeneous spatial distribution of components and irregular shapes of nucleoli, we selected the best-focused single slice for analyses. For the colocalization of endogenous NPM1, rRNA, SURF6, and FBL, we subdivided the nucleolus into several ROIs that encompassed the DFC and the inner GC regions. We chose these areas based on the established role of SURF6 as an early assembly factor for 60S pre-ribosomal subunits^15^. We reasoned that, since SURF6 binds to rRNA during the early maturation steps, focusing on regions surrounding the DFC and inner GC would give the highest probability to observe early assembly steps where NPM1 and SURF6 compete for rRNA binding. To establish the ROIs, we segmented the DFC regions of the nucleolus based on the intensities from the FBL channel and generated binary masks. We expanded the DFC masks using the dilation function of FIJI software. To segment the intensities for protein components (NPM1, SURF6, and FBL), we set a threshold to include only intensities above 25% of the maximum intensities. Due to the strong background and heterogeneous signals across samples for rRNA, we used thresholds at 15-40% of the maximum intensities for segmentation. We calculated the Mander’s overlap coefficient for ROIs identified in six individual nucleoli, each with multiple DFC regions.

To examine the effect of ActD treatment on the spatial distribution of the nucleolar components, we analyzed their colocalization within the entire nucleolus. To define the nucleolus, we segmented ROIs using the total intensity of NPM1, SURF6, rRNA, and FBL channels. We calculated the Pearson’s correlation coefficient for 40 DMSO-treated and 41 ActD-treated nucleoli.

For in vitro condensates, we calculated the Pearson’s correlation coefficient for entire condensates using masks generated using fluorescence intensities of the NPM1, SURF6, and rRNA channels.

### Recombinant protein expression and purification

Wild-type NPM1 (1-294) and its single cysteine mutant (C21T/C275T)^53^; and wild-type SURF6 (1-361) and its single cysteine mutant (C19S), each with an N-terminal 6X poly-histidine tag and a TEV recognition site sub-cloned into pET28 plasmids, were used for protein expression (Tables S2 and S3).

Wild-type NPM1 and its NPM1 (C21T/C275T) mutant were expressed and purified as previously described^53^. Briefly, bacterial cultures were grown up to an optical density (OD_600_) ∼ 0.8 at 37 °C, followed by overnight incubation at 18 °C post-induction with 0.4 mM IPTG. Cells were harvested via centrifugation and lysed in 25 mM Tris, 300 mM NaCl, 5 mM β-mercaptoethanol (BME), pH 7.5 by sonication. NPM1 was extracted from the soluble fraction via Ni-NTA affinity chromatography. His tags were cleaved with TEV protease during overnight dialysis at 4 °C in TBS-150 (10 mM Tris, 150 mM NaCl, 2 mM DTT, pH 7.5). The protein was further purified using a C4 HPLC column and pure NPM1 protein fractions were lyophilized. The lyophilized protein was re-suspended in a buffer consisting of 10 mM Tris (pH 7.5), 2 mM DTT, and 6 M GuHCl and subjected to refolding via dialysis against TBS-150 at 4 °C overnight.

BL21 (DE3) *E. coli* bacteria transformed with plasmids encoding wild-type SURF6 or SURF6-C19S were grown to O.D. ∼ 0.8 and induced for protein expression with 0.4 mM IPTG for 4 hours at 37 °C. Bacterial pellets were resuspended in 50 mM Tris, 300 mM NaCl, and 0.1% Triton X-100 pH 8.0 containing protease inhibitors (SigmaFast A, Millipore) and sonicated in ice. The pellet fraction was collected and resuspended in 50 mM sodium phosphate, 300 mM NaCl, and 5 mM BME at pH 7.5, 6 M GuHCl and homogenized again by sonication. The resuspension was centrifuged to separate undissolved cell debris, and the supernatant was passed through a Ni-NTA column in the presence of 6 M urea. SURF6-containing fractions were dialyzed at 4 °C in 20 mM Tris, 300 mM NaCl, pH 7.5, and 0.5 mM TCEP in the presence of TEV protease. The cleaved protein was further purified on a Mono-S column using a shallow salt gradient from 150 mM to 1 M NaCl in 20 mM Tris pH 7.5, 6 M urea, and 5 mM DTT. The protein-containing fraction was dialyzed in 10 mM Tris pH 7.5, 1 M NaCl, and 2 mM DTT at 4 °C overnight.

### Expression and purification of 70S ribosomes and total rRNA

The 70S ribosomes were purified from E. coli K12, strain A19, grown in Luria Broth. Cultures were grown at 37 °C to an OD_600_ ∼ 0.8 and harvested by centrifugation. Cell pellets were suspended in B-Per reagent (Thermo Fisher Scientific) supplemented with lysozyme and Turbo DNase (Thermo Fisher Scientific). The cell lysate was centrifuged at 30,000 g, 4 °C for 1 h. The soluble fraction was layered onto a 30% sucrose cushion prepared in 20 mM Tris-HCl, 50 mM MgOAc, 100 mM NH4Cl, 2 mM DTT, 0.5 mM EDTA, pH 7.5 buffer (buffer D) and centrifuged at 30,000 g, for 16 h at 4 °C in a SW 32 Ti Swinging-Bucket Rotor (Beckman Coulter). The resulting pellet was gently suspended in buffer D and subjected to two additional rounds of purification on a 30% sucrose cushion. The resulting pellet was dissolved in 10 mM Tris-HCl, 6 mM MgOAc, 50 mM NH4Cl, 1 mM TCEP, 0.5 mM EDTA, pH 7.5 buffer, and centrifuged at 16,000 g, 4 °C for 10 mins. The supernatant containing pure 70S ribosomes was aliquoted and stored at -80 °C. For phase separation assays, 70S ribosomes were dialyzed against TBS-150 containing 6 mM MgCl_2_ and further stabilized through the addition of 3 mM MgCl_2_. Total rRNA was extracted from the 70S ribosomes using the chloroform:isoamyl alcohol Trizol extraction protocol^54^, and rRNA purity was estimated by measuring the A260/A280 ratio. The samples with ratios ⁓2.0 were further analyzed on a 1.2% 1X TBE agarose gel for quality control. Purified total rRNA was stored at -80 °C. The rRNA stock was diluted into TBS-150 buffer for the phase separation assays.

### Fluorescence labeling

Fluorescently labeled NPM1 was prepared as previously reported^53^. Briefly, lyophilized NPM1 (C21T/C275T) was resuspended in a 25 mM Tris, 2 mM TCEP, pH 7.5 containing 6 M GuHCl and mixed with a two-fold molar excess of Alexa-488 maleimide fluorophore (Invitrogen). Unbound dye and unlabeled proteins were removed through HPLC purification. To ensure that each NPM1 pentamer contains only a single fluorophore, labeled and unlabeled NPM1 were refolded at a 1:10 ratio.

SURF6 (C19S) was labeled with Alexa-647 maleimide (Invitrogen) at 1:2 protein:dye ratio under denaturing conditions (10 mM Tris, 2 mM DTT, pH 7.5, 6 M GuHCl) overnight at 4 °C. Samples were dialyzed in TBS-1M at 4 °C. Any remaining unbound dye was removed by ultrafiltration using a 10 kDa cut-off filter (Amicon). Preparation and imaging of ternary and binary condensates To prepare the NPM1-SURF6-rRNA ternary condensates, we pre-made NPM1-rRNA condensates by mixing 2% AF488-NPM1 and rRNA in TBS-150 containing 2 µM SYTO 40 dye (for rRNA visualization) on a 16-well chambered slide (Grace Bio-Labs) coated with Sigmacote and 1% Pluronic F-127. NPM1-rRNA condensates were allowed to settle at the bottom for 40 minutes at room temperature. Following this, 2% labeled SURF6 was added to the NPM1 -rRNA condensates. Condensates were allowed to equilibrate for three hours prior to imaging. Images were collected on a Zeiss LSM 780 Observer.Z1 through a Plan Apochromat 63X/1.4 objective lens.

Shell and core phase fusion events and the SURF6 accumulation experiments were recorded on a 3i Marianas system configured with a Yokogawa CSU-W1 confocal scanning system using an alpha Plan Apochromat 100x/1.46 objective. For the SURF6 accumulation experiments, images for the NPM1 channel, in addition to the SURF6 channel, were recorded to demarcate the shell phase. Two-channel images of the condensates were collected every 30 seconds. To measure the increase in SURF6 intensities over time for the two phases, we defined 1-micron diameter ROIs and recorded the mean intensities that were contained within either the core or shell phases. To correct for photobleaching resulting from image acquisition, we measured the rate of photobleaching of multiphase condensates labeled with 2% SURF6. The photobleaching rate was extracted from single exponential fits of the curve ^19^.

Phase diagrams of the binary condensates were generated using 2% fluorescently labeled proteins to visualize NPM1 or SURF6 over the range of reported concentrations. For condensates containing rRNA, 2 µM SYTO 40 dye was added. Samples were incubated for two hours prior to imaging. Images were collected on the 3i Marianas system described above and were processed using Fiji image processing software ^55^.

### ANS fluorescence measurements

10 μM 8-Anilino-1-naphthalenesulfonic acid ammonium salt (ANS) (Sigma-Aldrich, USA) was added to binary condensates. For ternary condensates, ANS dye was mixed with NPM1 and rRNA, followed by a 40-minute incubation period. Subsequently, SURF6 was added to the preformed NPM1-rRNA condensate solution, followed by incubation at room temperature for approximately 2 h before imaging in CultureWell 16-well chambered slides to reach equilibrium conditions. All images were collected on a Zeiss LSM 780 Observer.Z1 through a Plan Apochromat 63X/1.4 objective lens. ANS fluorescence was excited with a 405 nm diode laser, and emitted light was collected at 420-624 nm. All the images were processed using Fiji image processing software^55^.

### Engulfment of cores by shells and contact angle measurements

Chambered slides (Grace BioLabs, Bend, OR) were treated with 1% Pluronic F-127 solution (Sigma-Aldrich, USA), to make the surface hydrophilic. To prepare the core mimics, 25 ng/μL of rRNA and 2.5 μM of SURF6 were mixed, and for the shell, 5 μM NPM1 and 4 μM of SURF6 were mixed. After 40 mins, the core mimics were added to the pre-formed shell mimics and analyzed using time-lapse imaging. Images were acquired on a Zeiss Lattice Light Sheet 7 microscope. Volumetric image stacks were captured using dithered square virtual lattice s (Sinc3 30x1000; approximately 30 µm in length and 1,000 nm in width) and stage scanning with 0.2 µm step sizes, resulting in 145 nm deskewed z-steps. Images were captured using a pair of Hammamatsu Orca Fusion sCMOS cameras, with the emission cube comprised of a 640 beam splitter, a 495-550 emission filter for AF488, and a long-pass 655 filter for AF647. Volumes were captured once every 6 seconds using 5 ms planar exposures. Images were saved in .czi file format and subsequently deskewed using the Zeiss Zen Blu2 3.6 Software. Deskewed lattice light sheet volumes were converted into Arrivis SIS files using the Arrivis 4 Software (Zeiss).3D reconstruction and subsequent images and movies were created and exported from Arrivis 4D software. For contact angle measurements, three binary condensates were prepared: NPM1 - SURF6 (10 μM each), NPM1-rRNA (10 μM NPM1 with 50 ng/μL rRNA), and SURF6-rRNA (10 μM SURF6 with 50 ng/μL rRNA). Deskewed lattice light sheet volume stacks were acquired and projected in XZ to obtain the shape profile. The contact angles, which are the angle between the line tangent to the condensates and the contact line interior of the condensate, were measured using Fiji image processing software^55^.

### Determination of volume fraction, Kp, and ΔΔG

We analyzed the volume fractions of the cores Vf_(core)_ within multiphase droplets from the volume of the core regions, V_(core),_ and the volume of the entire condensate V_(total)_ (equation 1). V_(core)_ was determined by creating surface masks from thresholding the mean intensities of the rRNA channel. Masks to determine V_(total)_ were created from thresholding the sum of mean intensities of NPM1, SURF6, and rRNA to ensure coverage of the entire volume of condensates. Vf _(core)_ values were determined for individual condensates.

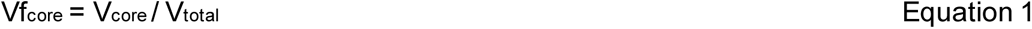

The apparent partition coefficient (K_app_) of NPM1 in homogeneous and multiphase condensates as a function of SURF6 concentrations was determined from mean intensities within the segmented masks from micrographs (Equations 2-4). For determining the K_app (dense)_ of homogeneous condensates, the NPM1 mean intensities inside homogenous condensates, I_dense_, were extracted from masks segmented from the sum of all channel intensities, which covered the entire condensate area. For the multiphase condensates, separate masks that represent the shell and core sub-phases of the condensates were generated. The mean intensities from the NPM1 (enriched component in the shell phases) channel were used to create the masks for the shell regions, while the rRNA (enriched component in the core phases) channel was used for the core regions. Following this, the NPM1 mean intensities (I_NPM1, shell_, I_NPM1, core_) were determined within the masks. To account for background signals, intensities from images of buffer solutions collected using identical imaging parameters were subtracted. To measure the light phase NPM1 intensities (I_NPM1 light_, the areas outside the condensates were masked, and the intensity threshold cut-off was manually adjusted when necessary to remove areas with out-of-focus condensates. Using the extracted mean intensities, the apparent partition coefficients, K_app_, were calculated from the NPM1 mean intensities (background subtracted), as previously reported^18^. All visualization and analyses of volumes and mean intensities were performed using IMARIS software.

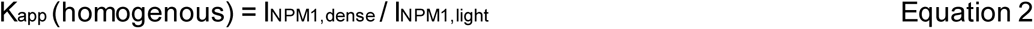

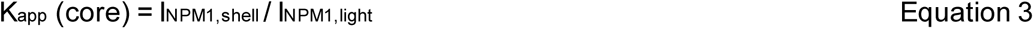

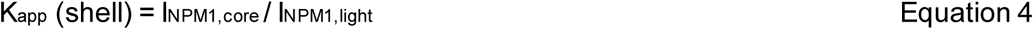

K_app_ values (Equation 5) of rRNA and 70S particles within NPM1-SURF6 condensates were calculated from the mean intensities of SYTO 40 dye from the masks of the entire condensate (I_SY TO 40, dense_) at 1:1 and 2:1 ratios of NPM1:SURF6 and the intensities from the light phase mask (I_SY TO 40, light)_). Transfer free energy (ΔG^tr^) values were calculated from the K_app_ values to evaluate the stability of rRNA or 70S interactions within the condensates (Equation 6). To determine the extent of destabilization of rRNA and 70S interactions, the change in rRNA transfer free energy was calculated as a function of increased NPM1 concentration, ΔΔG^tr^ (shown in Equation 7). ΔG^tr^_(2:1)_ and ΔG^tr^_(1:1)_ correspond to transfer free energies at 2:1 and 1:1 NPM1:SURF6 ratios, respectively. The change in transfer free energies was transformed to ΔΔG^Ef f lux^ (Equation 8) to represent the favorability for exclusion of the rRNA or 70S from condensates as interactions were weakened through the addition of excess NPM1.

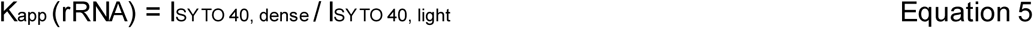

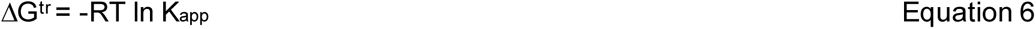

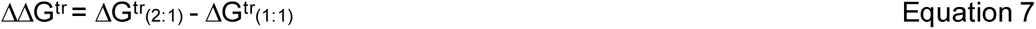

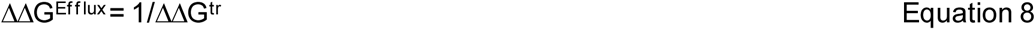

### Bioinformatics analyses of the pre-60S assembly factors

To identify unresolved regions in the cryo-EM structures of assembly factors in Homo sapiens^15,49^, we performed pairwise sequence alignments between the resolved sequences and their corresponding reference sequences from pre-60S assembly states A-H, encompassing a total of 36 assembly factors. Regions 30 amino acids long or longer were considered “resolved” for these analyses, and for each assembly factor, the state with the smallest and fewest resolved regions was retained for further analysis. Additionally, the unresolved regions, defined as those outside the resolved regions that contained segments shorter than 30 amino acids, were excluded from the analysis. The retained unresolved regions were analyzed using IUPRED2A (Mészáros B., et al., Nucleic Acids Res., 2018; Erdős G. and Dosztányi Z., Curr Protoc Bioinformatics, 2020, https://iupred2a.elte.hu/) to exclude ordered amino acid segments 30 amino acids long or longer with a disorder prediction score below 0.45. Finally, the fractions of arginine (Arg) and lysine (Lys) residues in the unresolved regions were calculated and compared to the human IDRome containing all IDRs in the human proteome. The human IDRome was generated from all human Swiss-Prot (reviewed) proteins contained in Uniprot release 2023_04. From this human proteome dataset, we identified 12,899 unique IDR sequences (from 8,067 proteins) using sak.stjude.org^56^ with lengths ≥ 60 residues, which is defined as the human IDRome.

## Supporting information

Video S1: Fusion events of shells and cores in multiphase condensates

Video S2: Engulfment of cores by pre-existing shells

Video S3: Accumulation of fl-SURF6 into the cores of multiphase condensates

## Acknowledgments

We thank Dr. Aaron Taylor from the Cell and Tissue Imaging Center-Light Microscopy for his support of fluorescence microscopy; the Center for Advanced Genome Engineering team at St. Jude Children’s Research Hospital for their contributions to genome engineering; the Vector Development and Production Shared Resource for providing lentiviral production; and the Hartwell Center team for Sanger DNA sequencing and STR profiling. Support for St. Jude core facilities is provided by the NCI Cancer Center Support Grant P30 CA021765 and by ALSAC. We also acknowledge funding from the St. Jude Research Collaborative on the Biophysics of RNP Granules (to R.W.K.) and NIH (R35GM131891 and R01GM115634 to R.W.K.).

## Author Contributions

P.D., M.C.F., and R.W.K. conceived the project. P.D., M.C.F., M.T., Q.M., S.M.P., S.T., and R.W.K. designed experiments. P.D., M.C.F., S.K., M.T., A.P., S.T., G.E.C., T.F., E.G., and C-G.P. performed the experiments and analyzed the data. P.D. and M.C.F. made the figures. P.D. wrote the first draft, and R.W.K. wrote/edited the manuscript. M.C.F., S.K., A.P., G.E.C., S.T., and S.M.P. contributed to writing and editing the Supplemental Information and Method Details of the manuscript. All authors discussed the results and provided feedback on the manuscript.

## References

1 Boeynaems, S. et al. Protein Phase Separation: A New Phase in Cell Biology. Trends Cell Biol 28, 420–435, doi:10.1016/j.tcb.2018.02.004 (2018).

2 Alberti, S. & Hyman, A. A. Biomolecular condensates at the nexus of cellular stress, protein aggregation disease and ageing. Nat Rev Mol Cell Biol 22, 196–213, doi:10.1038/s41580-020-00326-6 (2021).

3 Mitrea, D. M. & Kriwacki, R. W. Phase separation in biology; functional organization of a higher order. Cell Commun Signal 14, 1, doi:10.1186/s12964-015-0125-7 (2016).

4 Mittag, T. & Pappu, R. V. A conceptual framework for understanding phase separation and addressing open questions and challenges. Mol Cell 82, 2201–2214, doi:10.1016/j.molcel.2022.05.018 (2022).

5 Choi, J. M., Holehouse, A. S. & Pappu, R. V. Physical Principles Underlying the Complex Biology of Intracellular Phase Transitions. Annu Rev Biophys 49, 107–133, doi:10.1146/annurev-biophys-121219-081629 (2020).

6 Sabari, B. R., Dall’Agnese, A. & Young, R. A. Biomolecular Condensates in the Nucleus. Trends Biochem Sci 45, 961–977, doi:10.1016/j.tibs.2020.06.007 (2020).

7 Pederson, T. The nucleolus. Cold Spring Harb Perspect Biol 3, doi:10.1101/cshperspect.a000638 (2011).

8 Lafontaine, D. L. J., Riback, J. A., Bascetin, R. & Brangwynne, C. P. The nucleolus as a multiphase liquid condensate. Nat Rev Mol Cell Biol 22, 165–182, doi:10.1038/s41580-020-0272-6 (2021).

9 Correll, C. C., Bartek, J. & Dundr, M. The Nucleolus: A Multiphase Condensate Balancing Ribosome Synthesis and Translational Capacity in Health, Aging and Ribosomopathies. Cells 8, doi:10.3390/cells8080869 (2019).

10 Boisvert, F. M., van Koningsbruggen, S., Navascués, J. & Lamond, A. I. The multifunctional nucleolus. Nat Rev Mol Cell Biol 8, 574–585, doi:10.1038/nrm2184 (2007).

11 Leung, A. K. & Lamond, A. I. The dynamics of the nucleolus. Crit Rev Eukaryot Gene Expr 13, 39–54, doi:10.1615/critreveukaryotgeneexpr.v13.i1.40 (2003).

12 Tartakoff, A. et al. The dual nature of the nucleolus. Genes Dev 36, 765–769, doi:10.1101/gad.349748.122 (2022).

13 Feric, M. et al. Coexisting Liquid Phases Underlie Nucleolar Subcompartments. Cell 165, 1686–1697, doi:10.1016/j.cell.2016.04.047 (2016).

14 Kasciukovic, T. & Zomerdijk, J. C. B. M. in Encyclopedic Reference of Genomics and Proteomics in Molecular Medicine 1702–1707 (Springer Berlin Heidelberg, 2006).

15 Vanden Broeck, A. & Klinge, S. Principles of human pre-60S biogenesis. Science 381, eadh3892, doi:10.1126/science.adh3892 (2023).

16 Potapova, T. A. & Gerton, J. L. Ribosomal DNA and the nucleolus in the context of genome organization. Chromosome Research 27, 109–127, doi:10.1007/s10577-018-9600-5 (2019).

17 Mitrea, D. M. et al. Self-interaction of NPM1 modulates multiple mechanisms of liquid-liquid phase separation. Nat Commun 9, 842, doi:10.1038/s41467-018-03255-3 (2018).

18 Riback, J. A. et al. Composition-dependent thermodynamics of intracellular phase separation. Nature 581, 209–214, doi:10.1038/s41586-020-2256-2 (2020).

19 Ferrolino, M. C., Mitrea, D. M., Michael, J. R. & Kriwacki, R. W. Compositional adaptability in NPM1-SURF6 scaffolding networks enabled by dynamic switching of phase separation mechanisms. Nat Commun 9, 5064, doi:10.1038/s41467-018-07530-1 (2018).

20 King, M. R. et al. Macromolecular condensation organizes nucleolar sub-phases to set up a pH gradient. Cell 187, 1889–1906.e1824, doi:10.1016/j.cell.2024.02.029 (2024).

21 Mitrea, D. M. et al. Nucleophosmin integrates within the nucleolus via multi-modal interactions with proteins displaying R-rich linear motifs and rRNA. Elife 5, doi:10.7554/eLife.13571 (2016).

22 Tan, T. et al. Dynamic nucleolar phase separation influenced by non-canonical function of LIN28A instructs pluripotent stem cell fate decisions. Nat Commun 15, 1256, doi:10.1038/s41467-024-45451-4 (2024).

23 Quinodoz, S. A. et al. Mapping and engineering RNA-controlled architecture of the multiphase nucleolus. bioRxiv, 2024.2009.2028.615444, doi:10.1101/2024.09.28.615444 (2024).

24 Correll, C. C. et al. Crossing boundaries of light microscopy resolution discerns novel assemblies in the nucleolus. Histochem Cell Biol 162, 161–183, doi:10.1007/s00418-024-02297-7 (2024).

25 Sirri, V., Urcuqui-Inchima, S., Roussel, P. & Hernandez-Verdun, D. Nucleolus: the fascinating nuclear body. Histochem Cell Biol 129, 13–31, doi:10.1007/s00418-007-0359-6 (2008).

26 Hernandez-Verdun, D. The nucleolus: a model for the organization of nuclear functions. Histochem Cell Biol 126, 135–148, doi:10.1007/s00418-006-0212-3 (2006).

27 Politz, J. C., Polena, I., Trask, I., Bazett-Jones, D. P. & Pederson, T. A nonribosomal landscape in the nucleolus revealed by the stem cell protein nucleostemin. Mol Biol Cell 16, 3401–3410, doi:10.1091/mbc.e05-02-0106 (2005).

28 Politz, J. C., Lewandowski, L. B. & Pederson, T. Signal recognition particle RNA localization within the nucleolus differs from the classical sites of ribosome synthesis. J Cell Biol 159, 411–418, doi:10.1083/jcb.200208037 (2002).

29 Heintzmann, R. & Huser, T. Super-Resolution Structured Illumination Microscopy. Chem Rev 117, 13890–13908, doi:10.1021/acs.chemrev.7b00218 (2017).

30 Chen, X. et al. Superresolution structured illumination microscopy reconstruction algorithms: a review. Light Sci Appl 12, 172, doi:10.1038/s41377-023-01204-4 (2023).

31 Gibbs, E. et al. p14(ARF) forms meso-scale assemblies upon phase separation with NPM1. Res Sq, doi:10.21203/rs.3.rs-3592059/v1 (2023).

32 Jao, C. Y. & Salic, A. Exploring RNA transcription and turnover in vivo by using click chemistry. Proc Natl Acad Sci U S A 105, 15779–15784, doi:10.1073/pnas.0808480105 (2008).

33 Riback, J. A. et al. Viscoelasticity and advective flow of RNA underlies nucleolar form and function. Mol Cell 83, 3095–3107.e3099, doi:10.1016/j.molcel.2023.08.006 (2023).

34 Judy, E. & Kishore, N. A look back at the molten globule state of proteins: thermodynamic aspects. Biophysical Reviews 11, 365–375, doi:10.1007/s12551-019-00527-0 (2019).

35 Gussakovsky, E. E. & Haas, E. Two steps in the transition between the native and acid states of bovine alpha-lactalbumin detected by circular polarization of luminescence: evidence for a premolten globule state? Protein Sci 4, 2319–2326, doi:10.1002/pro.5560041109 (1995).

36 Matulis, D. & Lovrien, R. 1-Anilino-8-Naphthalene Sulfonate Anion-Protein Binding Depends Primarily on Ion Pair Formation. Biophysical Journal 74, 422–429, 10.1016/S0006-3495(98)77799-9 (1998).

37 Gasymov, O. K. & Glasgow, B. J. ANS fluorescence: potential to augment the identification of the external binding sites of proteins. Biochim Biophys Acta 1774, 403–411, doi:10.1016/j.bbapap.2007.01.002 (2007).

38 Geuskens, M. & Bernhard, W. [Ultrastructural cytochemistry of the nucleolus. 3. The effect of actinomycin D on the metabolism of nucleolar RNA]. Exp Cell Res 44, 579–598, doi:10.1016/0014-4827(66)90462-9 (1966).

39 Fu, Y. et al. Real-time imaging of RNA polymerase I activity in living human cells. J Cell Biol 222, doi:10.1083/jcb.202202110 (2023).

40 Iapalucci-Espinoza, S. & Franze-Fernández, M. T. Effect of protein synthesis inhibitors and low concentrations of actinomycin D on ribosomal RNA synthesis. FEBS Lett 107, 281–284, doi:10.1016/0014-5793(79)80390-7 (1979).

41 Shav-Tal, Y. et al. Dynamic sorting of nuclear components into distinct nucleolar caps during transcriptional inhibition. Mol Biol Cell 16, 2395–2413, doi:10.1091/mbc.e04-11-0992 (2005).

42 Maiser, A. et al. Super-resolution in situ analysis of active ribosomal DNA chromatin organization in the nucleolus. Scientific Reports 10, 7462, doi:10.1038/s41598-020-64589-x (2020).

43 Shan, L. et al. Nucleolar URB1 ensures 3′ ETS rRNA removal to prevent exosome surveillance. Nature 615, 526–534, doi:10.1038/s41586-023-05767-5 (2023).

44 Yip, S. P., Siu, P. M., Leung, P. H. M., Zhao, Y. & Yung, B. Y. M. The Multifunctional Nucleolar Protein Nucleophosmin/NPM/B23 and the Nucleoplasmin Family of Proteins. (The Nucleolus. 2011 May 23;15:213–52. doi: 10.1007/978-1-4614-0514-6_10. eCollection 2011.).

45 Ferrolino, M. C., Mitrea, D. M., Michael, J. R. & Kriwacki, R. W. Compositional adaptability in NPM1-SURF6 scaffolding networks enabled by dynamic switching of phase separation mechanisms. Nature Communications 9, 5064, doi:10.1038/s41467-018-07530-1 (2018).

46 Hong, Y. et al. Hydrophobicity of arginine leads to reentrant liquid-liquid phase separation behaviors of arginine-rich proteins. Nat Commun 13, 7326, doi:10.1038/s41467-022-35001-1 (2022).

47 Fisher, R. S. & Elbaum-Garfinkle, S. Tunable multiphase dynamics of arginine and lysine liquid condensates. Nat Commun 11, 4628, doi:10.1038/s41467-020-18224-y (2020).

48 Magoulas, C., Zatsepina, O. V., Jordan, P. W., Jordan, E. G. & Fried, M. The SURF-6 protein is a component of the nucleolar matrix and has a high binding capacity for nucleic acids in vitro. Eur J Cell Biol 75, 174–183, doi:10.1016/s0171-9335(98)80059-9 (1998).

49 Vanden Broeck, A. & Klinge, S. Eukaryotic Ribosome Assembly. Annual Review of Biochemistry 93, 189–210, 10.1146/annurev-biochem-030222-113611 (2024).

50 Cho, N. H. et al. OpenCell: Endogenous tagging for the cartography of human cellular organization. Science 375, eabi6983, doi:10.1126/science.abi6983 (2022).

51 Borer, R. A., Lehner, C. F., Eppenberger, H. M. & Nigg, E. A. Major nucleolar proteins shuttle between nucleus and cytoplasm. Cell 56, 379–390, doi:10.1016/0092-8674(89)90241-9 (1989).

52 Gibbs, E. et al. p14(ARF) forms meso-scale assemblies upon phase separation with NPM1. Nat Commun 15, 9531, doi:10.1038/s41467-024-53904-z (2024).

53 White, M. R. et al. C9orf72 Poly(PR) Dipeptide Repeats Disturb Biomolecular Phase Separation and Disrupt Nucleolar Function. Molecular Cell 74, 713–728.e716, doi:10.1016/j.molcel.2019.03.019 (2019).

54 Chomczynski, P. & Sacchi, N. The single-step method of RNA isolation by acid guanidinium thiocyanate–phenol–chloroform extraction: twenty-something years on. Nature Protocols 1, 581–585, doi:10.1038/nprot.2006.83 (2006).

55 Schindelin, J. et al. Fiji: an open-source platform for biological-image analysis. Nature Methods 9, 676–682, doi:10.1038/nmeth.2019 (2012).

56 Tripathi, S. et al. Defining the condensate landscape of fusion oncoproteins. Nat Commun 14, 6008, doi:10.1038/s41467-023-41655-2 (2023).

