## Supplemental Figures S1-S8 and Supplemental Tables S1-S3 for "Granular component sub-phases direct ribosome biogenesis in the nucleolus"

[¶]Contributed equally

[2]Current address: ElevateBio, 200 Smith Street, Waltham, MA 02451.

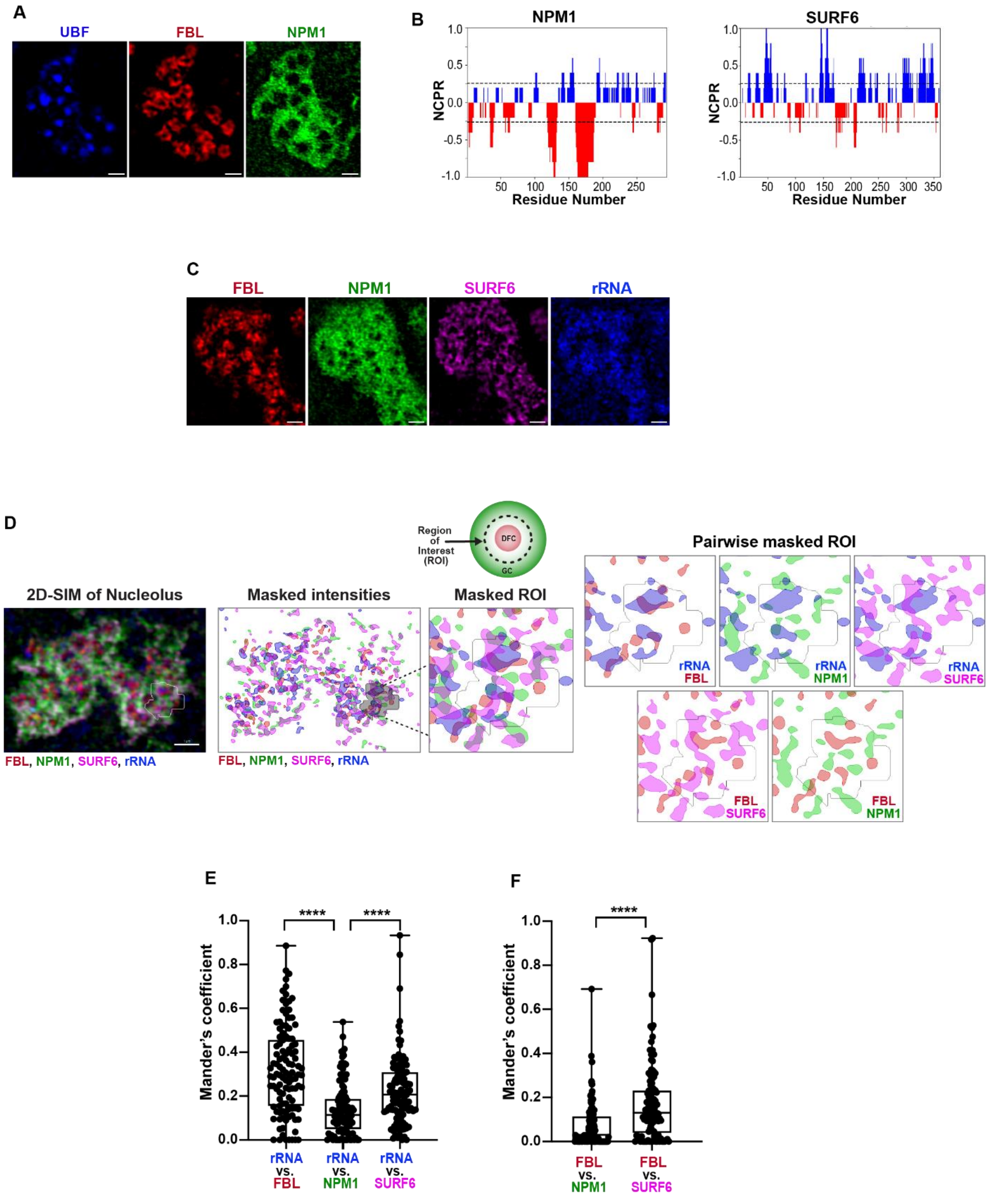


**Figure S1. Nucleolar components are heterogeneously distributed in the granular component.** (A) SIM image of a nucleolus immunostained for UBF (left), FBL-mCherry (center), and NPM1-mEGFP (right), scale bar = 1 µm. (B) Net charge per residue (NCPR) plots for NPM1

and SURF6 generated using CIDER analysis (Holehouse A., *et al., Biophys. J.,* 2017, https://pappulab.wustl .edu/CIDER/analysis/) illustrating the enrichment of acidic (Asp and Glu, red) and basic (Arg and Lys, blue) residues in NPM1 and SURF6. (C) SIM images of a nucleolus immunostained for FBL (left), NPM1 (center, left), SURF6 (center, right), and rRNA (right), scale bar = 1 µm. See Methods Details. (D) Segmentation of ROIs analyzed to determine colocalization of FBL, NPM1, SURF6, and rRNA. Shown are the overlaid images of a nucleolus for NPM1, rRNA, SURF6, and FBL channels (left) and the masks generated from segmentation of each component (labeled "Masked intensities and "Masked ROI"), scale bar = 1 µm. ROIs containing the DFC and inner GC regions were defined using the FBL mask, which was expanded using the Fiji dilation function. An example ROI is indicated by the white or gray outlines. The panels on the right (labeled "Pairwise masked ROI") show masks for pairs of components enclosed by an ROI. This type of ROI was used to compute Pearson correlation coefficient values for pairs of fluorescently labeled components. (E) Mander's overlap coefficient values for rRNA with FBL, NPM1, and SURF6 (n = 84 ROIs). (F) Mander's overlap coefficient values for FBL with NPM1 and SURF6, showing greater colocalization of FBL with SURF6 than with NPM1 (n = 112 ROIs).

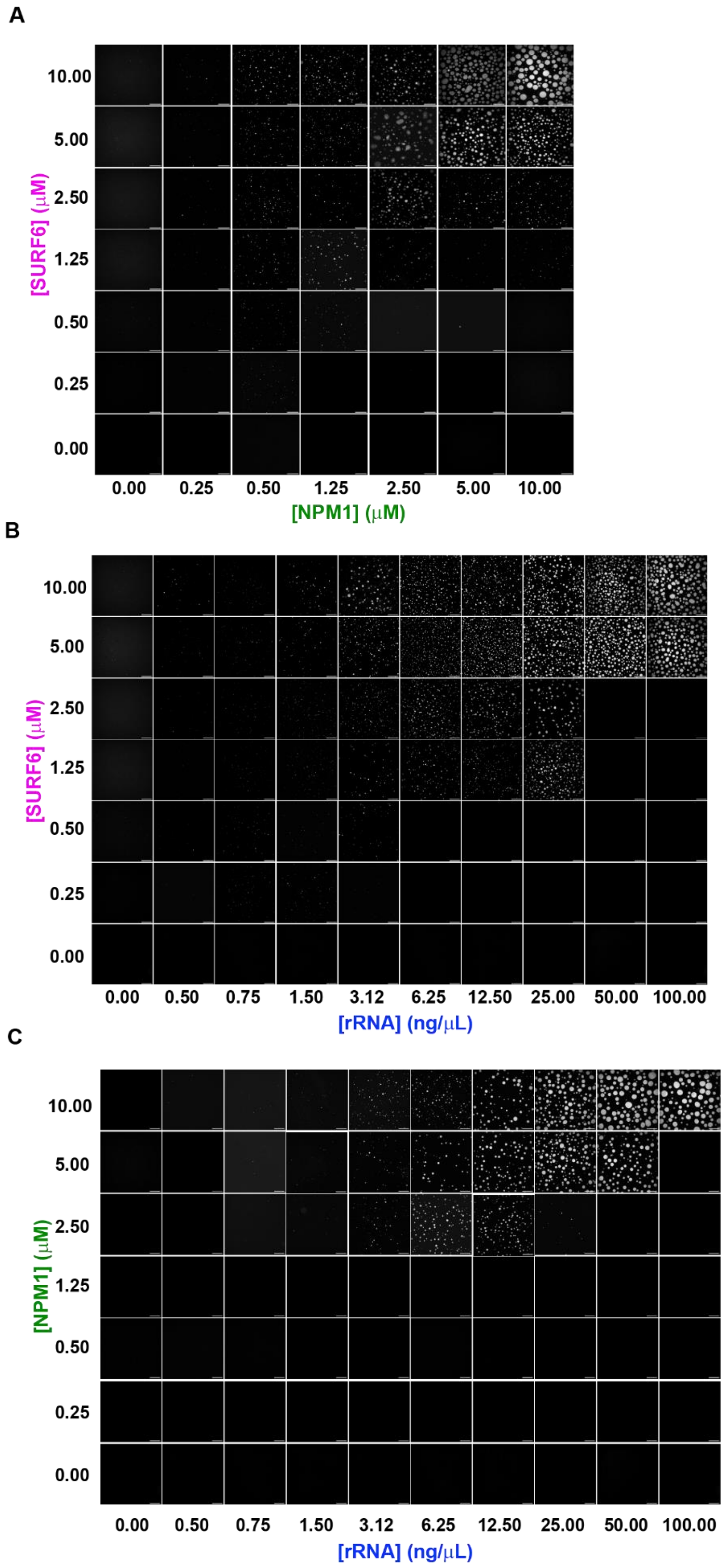
A
[SURF6] (μM)
10.00
5.00
2.50
1.25
0.50
0.25
0.00
0.00 0.25 0.50 1.25 2.50 5.00 10.00
[NPM1] (μM)
B
[SURF6] (μM)
10.00
5.00
2.50
1.25
0.50
0.25
0.00
0.00 0.50 0.75 1.50 3.12 6.25 12.50 25.00 50.00 100.00
[rRNA] (ng/μL)
C
[NPM1] (μM)
10.00
5.00
2.50
1.25
0.50
0.25
0.00
0.00 0.50 0.75 1.50 3.12 6.25 12.50 25.00 50.00 100.00
[rRNA] (ng/μL)

**Figure S2. Analysis of condensates in single-component and two-component mixtures of NPM1, SURF6, and rRNA.** Overlaid fluorescence micrographs (represented in grayscale) of single component and binary mixtures of NPM1 and SURF6, SURF6 and rRNA, and NPM1 and rRNA (A-C) in 10 mM Tris, 150 mM NaCl, 2 mM DTT pH 7.5. NPM1 and SURF6 were conjugated with AF488 and AF647, respectively. rRNA was labeled with SYTO 40 dye. Fluorescence intensities for the relevant labeled species were combined to give the grayscales, scale bar = 10 μm.

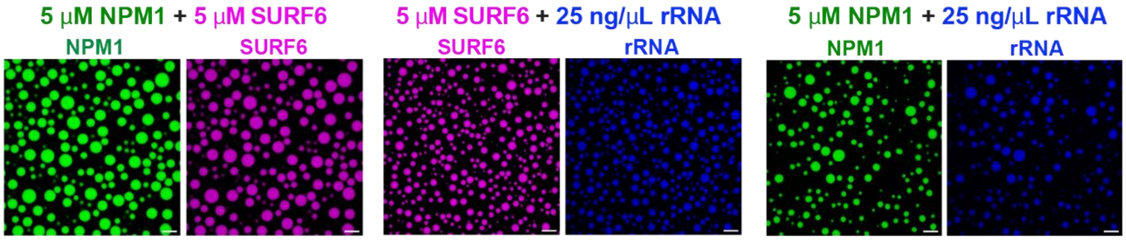


**Figure S3. Fluorescence images of condensates formed by binary mixtures of NPM1, SURF6, and rRNA.** Fluorescence micrographs of binary mixtures of NPM1 (green) and SURF6 (magenta) (left pair), SURF6 (magenta) and rRNA (blue) (center pair), and NPM1 and rRNA (right pair) at the noted concentrations in 10 mM Tris, 150 mM NaCl, 2 mM DTT pH 7.5. NPM1 (green) and SURF6 (magenta) were conjugated with AF488 and AF647, respectively, and rRNA (blue) was labeled with SYTO 40 dye, scale bar = 10 μm.

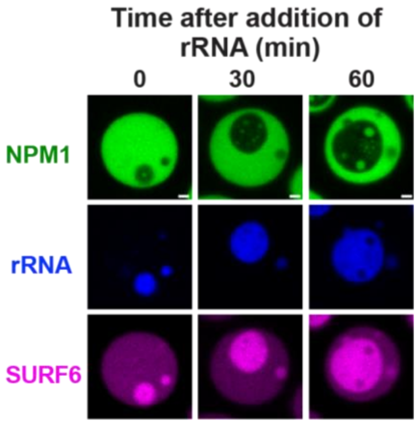


**Figure S4. Dependence of core sub-phase volume fraction on rRNA concentration.** Fluorescence micrographs of a NPM1-SURF6-rRNA condensate show an increase in the size of the core sub-phase over time after the addition of a three-fold excess (18.75 ng/μL) of rRNA. NPM1 (green) and SURF6 (magenta) were conjugated with AF488 and AF647, respectively, and rRNA (blue) was labeled with SYTO 40 dye, scale bar = 1 μm. The condensates initially contained 5 μM NPM1, 5 μM SURF6, and 6.25 ng/μL rRNA.

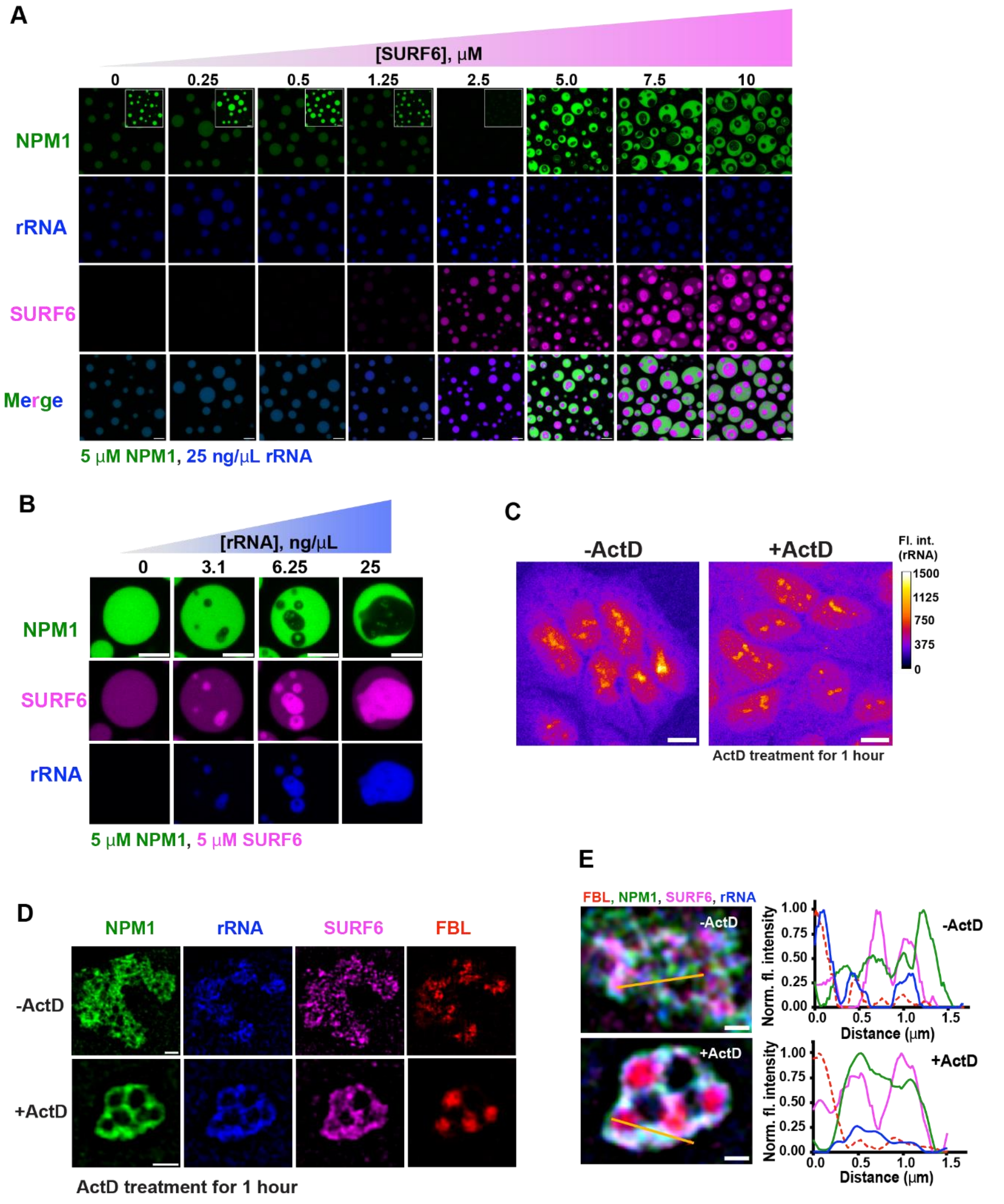


**Figure S5. Component concentration dependence of multiphase condensate formation and spatial heterogeneity of the GC.** (A) Fluorescence micrographs of NPM1-rRNA condensates titrated with SURF6, scale bar = 5 μm . NPM1 is shown in green (top row), rRNA in blue (second row), and SURF6 in magenta (third row). The bottom row shows an overlay of the three individual channels. Condensates contain 5 μM NPM1 and 25 ng/μL rRNA and the indicated concentrations of SURF6. The fluorescence intensity threshold for the inset images in the top row was reduced 4-fold. (B) Fluorescence micrographs of NPM1-SURF6 condensates show increased core sub-phase volume with increased rRNA concentrations, scale bar = 5 μm. NPM1 is shown in green (top row), SURF6 in magenta (middle row), and rRNA in blue (bottom row). Condensates contain 5 μM NPM1 and 5 μM SURF6 and the indicated concentrations of rRNA. (C) Fluorescence micrographs (shown in fire LUT) of ActD untreated (left) and treated (right) DLD-1$^{NPM1\text{-}G/FBL\text{-}R}$ cells labeled with EU-Alexa-647, scale bar = 1 μm. (D) SIM images of individual channels for nucleoli from ActD untreated (top panels) and treated (bottom panels) DLD-1$^{NPM1\text{-}G/FBL\text{-}R}$ cells. NPM1 is shown in green (first column), rRNA in blue (second column), SURF6 in magenta (third column), and FBL is shown in red (fourth column), scale bar = 1 μm. (E) Overlaid channels of SIM images of nucleoli with labeled FBL (red), NPM1 (green), SURF6 (magenta), and rRNA (blue) from ActD untreated (top left) and treated (bottom left) DLD-1$^{NPM1\text{-}G/FBL\text{-}R}$ cells, scale bar = 1 μm. The panels on the right show representative line profile plots (indicated by orange line in the images) for the four channels in ActD untreated (top) and treated (bottom) nucleoli.

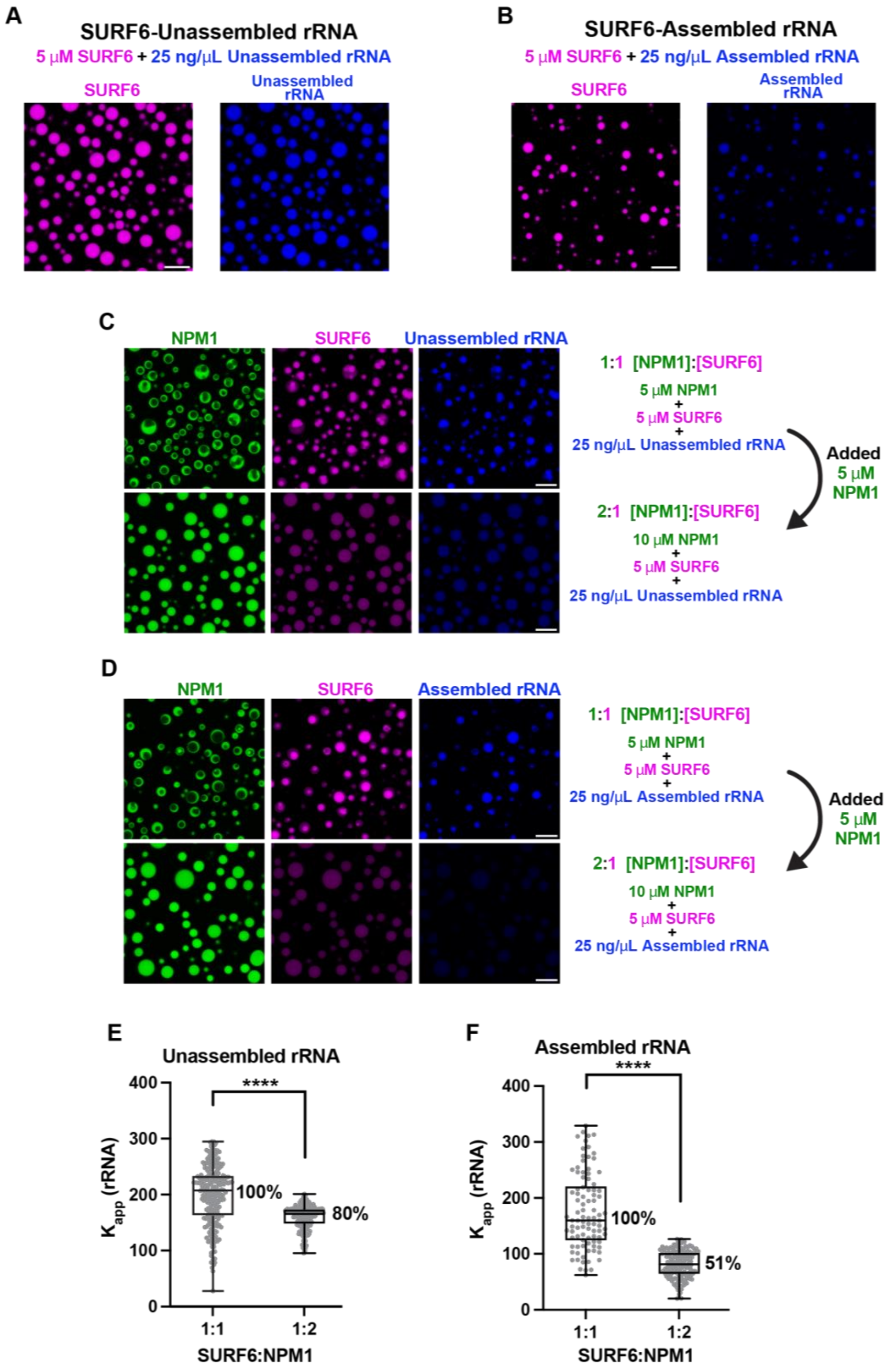

A
SURF6-Unassembled rRNA
5 μM SURF6 + 25 ng/μL Unassembled rRNA
SURF6
Unassembled rRNA
B
SURF6-Assembled rRNA
5 μM SURF6 + 25 ng/μL Assembled rRNA
SURF6
Assembled rRNA
C
NPM1
SURF6
Unassembled rRNA
1:1 [NPM1]:[SURF6]
5 μM NPM1 + 5 μM SURF6 + 25 ng/μL Unassembled rRNA
Added 5 μM NPM1
2:1 [NPM1]:[SURF6]
10 μM NPM1 + 5 μM SURF6 + 25 ng/μL Unassembled rRNA
D
NPM1
SURF6
Assembled rRNA
1:1 [NPM1]:[SURF6]
5 μM NPM1 + 5 μM SURF6 + 25 ng/μL Assembled rRNA
Added 5 μM NPM1
2:1 [NPM1]:[SURF6]
10 μM NPM1 + 5 μM SURF6 + 25 ng/μL Assembled rRNA
E
Unassembled rRNA
****
K app (rRNA)
0
100
200
300
400
100%
80%
1:1
1:2
SURF6:NPM1
F
Assembled rRNA
****
K app (rRNA)
0
100
200
300
400
100%
51%
1:1
1:2
SURF6:NPM1

**Figure S6. Weakened SURF6-rRNA interactions lead to the exclusion of assembled rRNA from SURF6-NPM1 condensates.** (A) Fluorescence micrograph of individual channels for condensates made with SURF6 (left, magenta) and unassembled rRNA (right, blue), scale bar = 10 μm. (B) Fluorescence micrographs of individual channels for condensates made with SURF6 (left, magenta) and assembled rRNA (right, blue), scale bar = 10 μm. SURF6 is conjugated with AF647, and unassembled/assembled rRNA was stained with SYTO 40 dye in A and B. (C) Fluorescence micrographs of individual channels for condensates made with SURF6 (center, magenta), unassembled rRNA (right, blue), and 5 μM (top row) or 10 μM (bottom row) NPM1 (left, green), scale bar = 10 μm. (D) Fluorescence micrographs of individual channels for condensates made with SURF6 (center, magenta), assembled rRNA (right, blue), and 5 μM (top row) or 10 μM (bottom row) NPM1 (left, green), scale bar = 10 μm. NPM1 and SURF6 were conjugated with AF488 and AF647, respectively; unassembled/assembled rRNA was stained with SYTO 40 dye in C and D. (E) Plot of partition coefficient ($K_{app}$) values for unassembled rRNA in SURF6:NPM1 condensates at 1:1 and 1:2 ratios of SURF6 and NPM1. (F) Plot of partition coefficient ($K_{app}$) values for assembled rRNA in SURF6:NPM1 condensates at 1:1 and 1:2 ratios of SURF6 and NPM1.

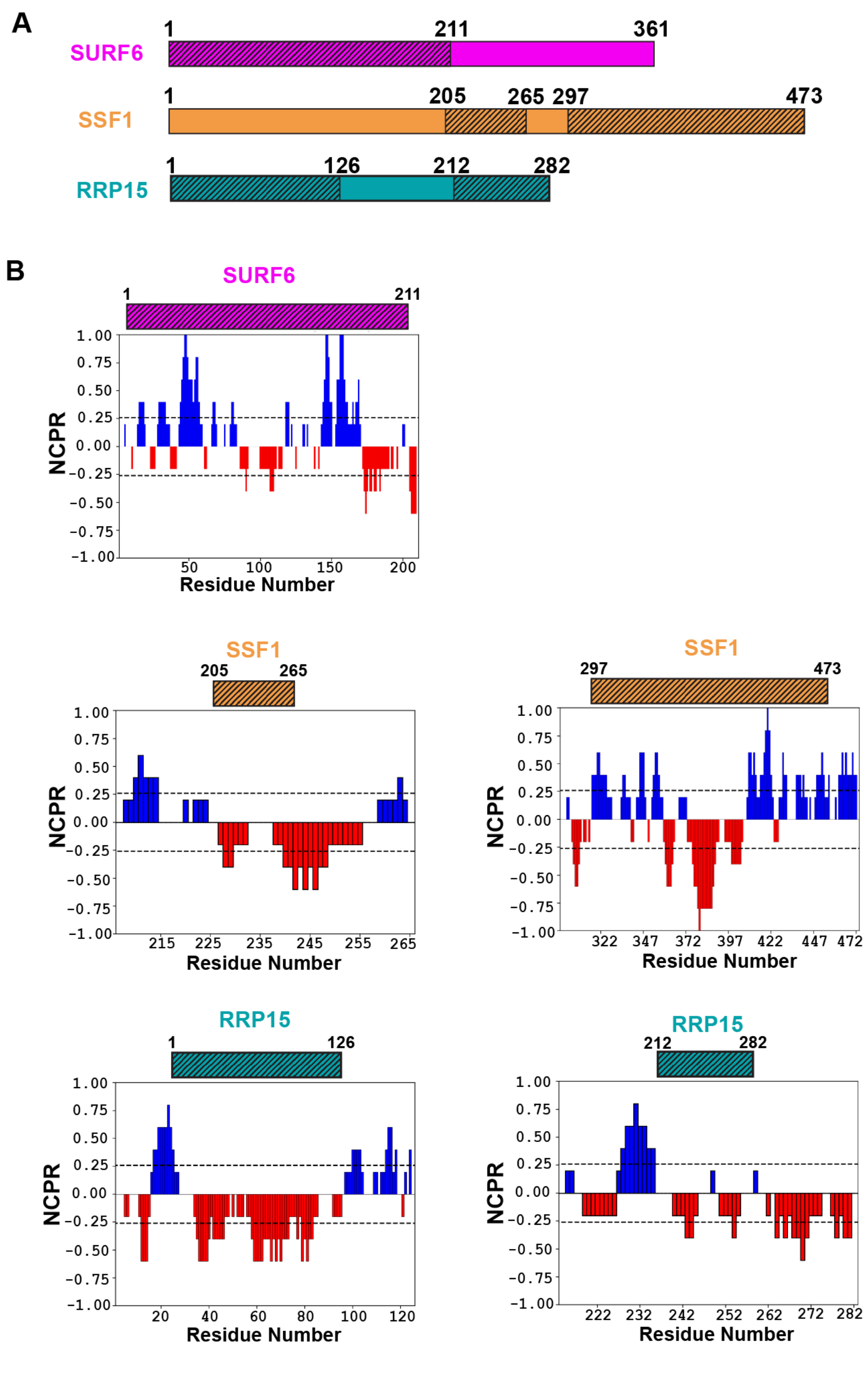


**Figure S7. Structurally unresolved regions of SURF6, SSF1, and RRP15 are enriched in charged residues.** (A) Schematic representation of the unresolved regions (indicated as hashed bars) present in SURF6 (1-211), SSF1 (205-265 and 297-473), and RRP15 (1-126 and

212-282). The unresolved regions were obtained from pre-60S ribosomal subunit assembly States A and B (Vanden Broeck A, Klinge S., *Science*. 2023; see Methods Details). (B) Net charge per residue (NCPR) plots for unresolved regions of SURF6, RRP15 and SSF1 from panel a generated using CIDER analysis (Holehouse A., *et al., Biophys. J.,* 2017, https://pappulab.wustl .edu/CIDER/analysis/) illustrating enrichment of acidic (Asp and Glu, red) and basic (Arg and Lys, blue) residues.

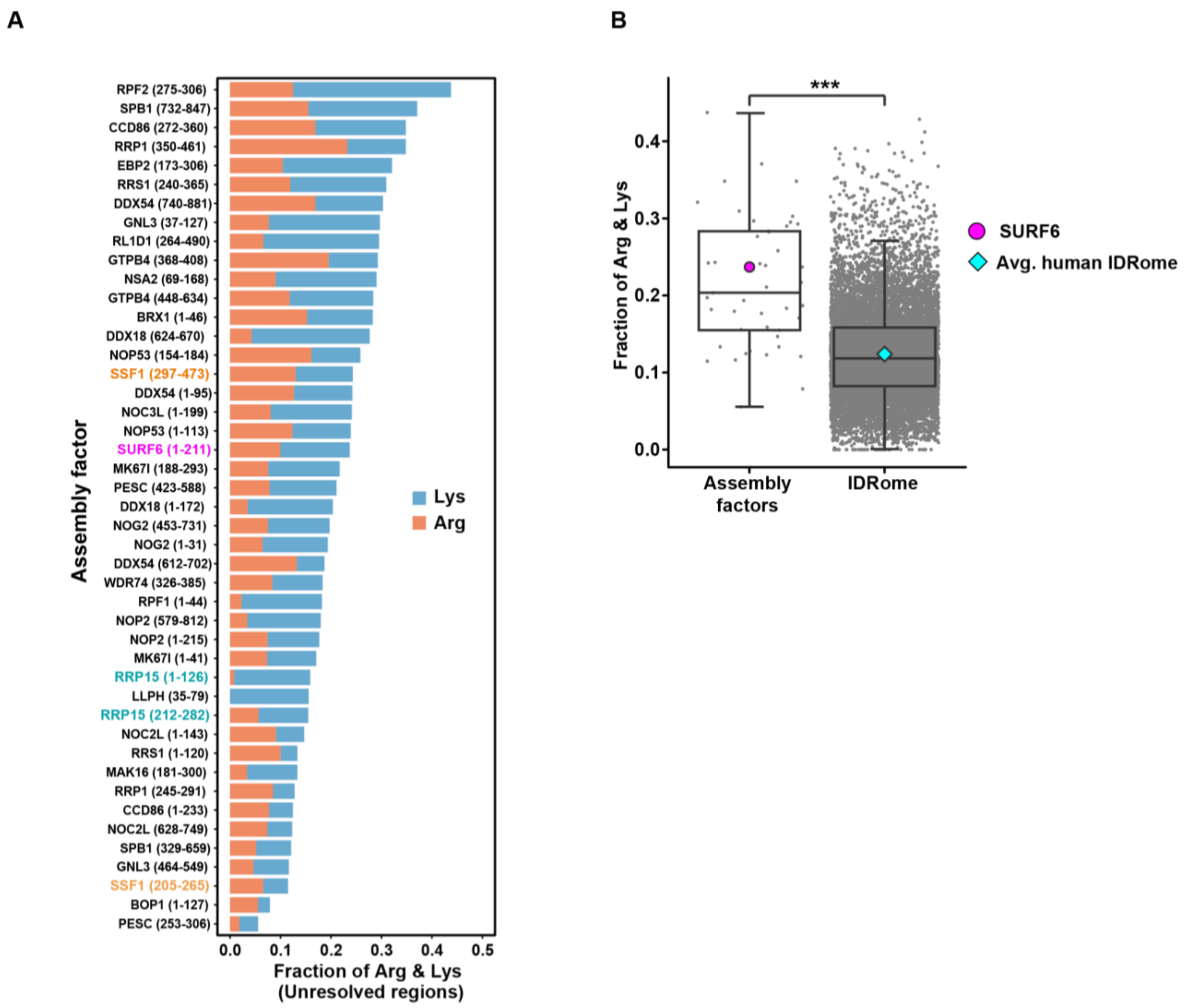


**Figure S8. Structurally unresolved regions of pre-60S assembly factors are enriched in Arg and Lys residues.** (A) Bar plot of the fraction of Arg and Lys residues in the unresolved regions of pre-60S assembly factors from 12 structurally characterized assembly states (A-H) (Vanden Broeck A, Klinge S., *Science*. 2023; see Methods Details). (B) Box plot of fractions of Arg and Lys in the unresolved regions of the assembly factors (left) shown in panel A. The fraction of Arg and Lys residues in the unresolved region of SURF6 (left) is shown as a magenta circle. Box plot of fractions of Arg and Lys for the human IDRome (right); the average value is shown as a blue diamond. Number of unresolved regions (n) = 45, and the number of IDRs in the human IDRome (n) = 12,899. The p-value was calculated using the Welch two-sample t-test

(***, $p < 0.001$) by comparing the fraction of Arg and Lys in the unresolved regions and the human IDRome.

### Supplemental Tables

**Table S1: Edited construct sequences and screening primers to generate DLD1-NPM1-mEGFP/FBL-mCherry C-terminally double-tagged cells.**

| Name | Sequence (5' to 3') |
|---|---|
| CAGE117.NPM1.g1 spacer | UCCAGGCUAUUCAAGAUCUC |
| CAGE117.g1.mEGFP donor (homology arms and insert)<br><br>Homology arms in lowercase<br><br>mEGFP in uppercase<br><br>5X GGS linker in bold uppercase<br><br>Silent blocking mutations in italics uppercase | ctcaggtgatccaacaccttggcctcttaaagtgctgggattacaggcatgagcc<br>accatgcctggccagctgttttttttgttggtttgttttttgttttggtacccatctgtagtgt<br>gatcttggctcactgcaacctctgcctcttgggctcaggcagtcctcccacctcag<br>cctcctgagtagctgggcctcctgtagttgcacaccaccaagcctggctaattttg<br>catttttagtagacagggtttcaccatgttgcccaggctggtctcaaattcctgagct<br>gaagtgatctgcccgcctcagtctcccaaagtgtagggattacaggcgtgagcc<br>accatgcctagcctcagcatatagtttttctaaatgtacacatgcccaggcacac<br>atgcacaggcaattcagaataagtttctggtgtttatgtaactttatttgccaaatctg<br>gccaactctaaagctgatctcgggagatgaagttggaagtaacattggccatat<br>gggtctctgttctttctgttgatttccttaagtaaataatgctaaactattaaataattatt<br>agtatattgttcacattttatgactgattaaagtgtttggaattaaattacatctgagt<br>ataaattttcttggagtcatatctttatctagagttaactctctggtggtagaatgaaa<br>aatagatgttgaactatgcaaagagacatttaatttattgatgtctatgaagtgttgt<br>ggttccttaaccacatttctttttttttttttccaggctattcaaga*C*ct*G*tggca*A*tgg<br>*C*g*A*aa*AAGC*ct*G***GGAGGAAGCGGAGGTTCTGGCGGTA**<br>**GTGGTGGATCTGGCGGCAGC**ATGGTTTCCAAGGGCG<br>AAGAACTGTTCACCGGCGTGGTGCCCATTCTGGTGGA<br>ACTGGACGGGGATGTGAACGGCCACAAGTTTAGCGTT<br>AGCGGCGAAGGCGAAGGGGATGCCACATACGGAAAG<br>CTGACCCTGAAGTTCATCTGCACCACCGGCAAGCTGC<br>CTGTGCCTTGGCCTACACTGGTCACCACACTGACATAC<br>GGCGTGCAGTGCTTCAGCAGATACCCCGACCATATGA<br>AGCAGCACGACTTCTTCAAGAGCGCCATGCCTGAGGG<br>CTACGTGCAAGAGCGGACCATCTTCTTTAAGGACGAC<br>GGCAACTACAAGACCAGGGCCGAAGTGAAGTTCGAGG<br>GCGACACCCTGGTCAACCGGATCGAGCTGAAGGGCAT<br>CGACTTCAAAGAGGACGGCAACATCCTGGGCCACAAG<br>CTCGAGTACAACTACAACAGCCACAACGTGTACATCAT<br>GGCCGACAAGCAGAAAAACGGCATCAAAGTGAACTTC<br>AAGATCCGGCACAACATCGAGGACGGCTCTGTGCAGC<br>TGGCCGATCACTACCAGCAGAACACACCCATCGGAGA<br>TGGCCCTGTGCTGCTGCCCGATAACCACTACCTGAGC<br>ACCCAGAGCAAGCTGAGCAAGGACCCCAACGAGAAGC<br>GGGACCACATGGTGCTGCTGGAATTTGTGACAGCCGC<br>CGGAATCACCCTCGGCATGGATGAGCTGTACAAGTAA<br>gaaaatagtttaaacaatttgttaaaaaattttccgtcttatttcatttctgtaacagttg<br>atatctggctgtcctttttataatgcagagtgagaactttccctaccgtgtttgataaat<br>gttgtccaggttctattgccaagaatgtgttgtccaaaatgcctgtttagtttttaaag<br>atggaactccaccctttgcttggttttaagtatgtatggaatgttatgataggacata<br>gtagtagcggtggtcagacatggaaatggtggggagacaaaaatatacatgtg |

| | |
|---|---|
| | aaataaaactcagtattttaataaagtagcacggtttctattgacttatttaactgcttt<br>atactttgtcaaagaaataattaatgtagttaggaatggcaaatagtcttgtaaaat<br>tctatgagaatgtccctgccctccccttcaatattctctctggagctaaccactttttca<br>tcataaggatttagtgctgtgttcccacctcctgatgatagttaacaattattataact<br>atgcaacatgtttccaaatgttccattagacctcctatctgcctattctagcctcactt<br>gcaaagaaaatgtggcatgttaaaacagcttaaaagcagcctttcaacctgtat<br>ggtttttcccctaggctggagtgcagtggcacaatctcagcttattgcagcttctgct<br>tcttgggttcaagcaggtctcctgcctcagcctcccaagtagctgggattacaggt<br>gtgagccaccagcccggctaatttttgtatttttagtagaga |
| CAGE117.gen.F2 | TGTACCTGAGAACCCATTGGC |
| CAGE117.junc.mEGFP.DS.R2 | GTTCACATCCCCGTCCAGTT |
| CAGE117.junc.mEGFP.DS.F2 | GCTGCCCGATAACCACTACC |
| CAGE117.gen.R2 | AGGCAGAACATATAAAGGTGCTAAT |
| CAGE117.DS.F | AGTTAACTCTCTGGTGGTAGAATGA |
| CAGE117.DS.R | CCAAGCAAAGGGTGGAGTTC |
| CAGE223.FBL.g3 spacer | AACUGAAGUUCAGCGCUGUC |
| CAGE223.g3.mCherry.donor<br><br>(homology arms and insert)<br><br>Homology arms in lowercase<br><br>mCherry in uppercase<br><br>5X GGS linker in bold uppercase | caccgtgttggccaggatggtctcaatctcctgatttcgtgattgagccacctcggc<br>ctcccaaagtgctgggattacaggcgtgagccaccacgcccagccttagactg<br>ggtaatttataatgaatggaaatttatttggctcccagttccaaaggctggaaagtc<br>caagattggaggtctgaatctggcgagggccttcttgctgtcatccattggcagaa<br>gggtgagagcaagatagaaagggggcataatcatccttttaatcagcaaccca<br>ctcttgtgataatagcattactctattcaggaaggcagaggcctcatgacctgaat<br>catctctcgaaggtcccacctctcaactcttgcatttaagggttacgtttccaacac<br>atgaactttggggggacacactagaaccatagcactgagttttacttgaattaataa<br>tgaaaacatctggtttaaagagcacacaagagaaaaacagcccaaagccctg<br>ttgtagacattagtcctttctcctctttaggccaactgcattgactccacagcctcag<br>ccgaggccgtgtttgcctccgaagtgaaaaagatgcaacaggagaacatgaa<br>gccgcaggagcagttgacccttgagccatatgaaagagaccatgccgtggtcg<br>tgggagtgtacaggtgagcaggggcccagcaatacaccaagacagacatctc<br>tgtcccttgcaccccgagtgccatgatcctggggaccctccttcatcacctatcttc<br>ctctcacaggccacccccaaggtgaagaac**GGAGGAAGCGGAGG**<br>**TTCTGGCGGTAGTGGTGGATCTGGCGGCAGC**ATGGT<br>GTCTAAGGGCGAAGAGGACAACATGGCCATCATCAAA<br>GAATTCATGCGGTTCAAGGTGCACATGGAAGGCAGCG<br>TGAACGGCCACGAGTTCGAGATTGAAGGCGAAGGCGA<br>GGGCAGACCTTACGAGGGAACACAGACCGCCAAGCT<br>GAAAGTGACCAAAGGCGGCCCTCTGCCTTTTGCCTGG<br>GACATTCTGAGCCCTCAGTTTATGTACGGCAGCAAGG<br>CCTACGTGAAGCACCCCGCCGATATTCCCGACTACCT<br>GAAGCTGAGCTTCCCCGAGGGCTTCAAGTGGGAGAGA<br>GTGATGAACTTCGAGGACGGCGGCGTGGTCACCGTGA<br>CTCAAGATAGCTCTCTGCAGGACGGCGAGTTCATCTA<br>CAAAGTGAAGCTGCGGGGCACCAACTTTCCCTCTGAT<br>GGCCCCGTGATGCAGAAAAAGACCATGGGCTGGGAA<br>GCCAGCAGCGAGAGAATGTACCCTGAAGATGGCGCCC<br>TGAAGGGCGAGATCAAGCAGCGGCTGAAACTGAAGGA<br>TGGCGGCCACTACGACGCCGAAGTGAAAACCACCTAC<br>AAGGCCAAGAAACCCGTGCAGCTGCCTGGCGCCTACA<br>ACGTGAACATCAAGCTGGACATCACCAGCCACAACGA<br>GGACTACACCATCGTGGAACAGTACGAGAGAGCCGAA |

| | |
|---|---|
| | GGCAGACACAGCACAGGCGGAATGGACGAGCTGTAC<br>AAGtgaagttcagcgctgtcaggattgcgagagatgtgtgttgatactgttgcac<br>gtgtgtttttctattaaaagactcatccgtctcccatgtctgctgctcattcctcccc ttg<br>acctgctgacacagggagcacgcacccttggtcaattttgcggggttgggtaaat<br>tctcactcggtcacagagcgcatgctccgtttctagctgcctttgcgcagcggcag<br>cctggatttcggttcttgggtgggattggtagctcgctgcgcatgcgtgcaggtaag<br>cggccatctcgcgcaggcggagtgtcagtgtgggtcacgtgaggggagcgga<br>gagggagggatgggggcggagtccagggcgtgggggggccggtttgttgtggt<br>cgccattttgctggttgcattactgggtaatcggggccctggcttgccgcgtccgcc<br>ggataccctcagccagtgggcaggtctgagctcgggctccccgagcagtttgag<br>tccccttgcccgctccttcaggtaacggcgcggggacgggtggggcggcaagc<br>ggtcgcagggaggtgggcaggacgggatccgccctgctcccgtcgccgtgag<br>acttagcacgaggccaagggaggagaggagggggggtggcaggcaggtgcg<br>ggccctgcctggctattcatagttgaattcctggaaccggccaagcccgaggaa<br>gcagttgcaggagggaggctgggaggggggtagccgggccccactcccgccct<br>ttgtttgggctcagctccgcgggccgcttcttcgtcgcctagcaacagctgcccta |
| CAGE223.FBL.DS.F | GCCGTGGTCGTGGGAGTGTACAGGT |
| CAGE223.FBL.DS.R | AATTGACCAAGGGTGCGTGCTCCC |
| CAGE223.gen.F | GCCTTTTATCACATTTCCTAAACCC |
| CAGE223.junc.DS.R | CATGTTGTCCTCTTCGCCCT |
| CAGE223.junc.DS.F | TGGACATCACCAGCCACAAC |
| CAGE223.gen.R | ACAACTGTACCATTGGTCGCT |

**Table S2: Amino acid sequence of NPM1 and SURF6.**

| Construct Name | Amino Acid Sequence |
| --- | --- |
| Wild-type NPM1, in vitro | GSMEDSMDMDMSPLRPQNYLFGCELKADKDYHFKVDN<br>DENEHQLSLRTVSLGAGAKDELHIVEAEAMNYEGSPIKVT<br>LATLKMSVQPTVSLGGFEITPPVVLRLKCGSGPVHISGQH<br>LVAVEEDAESEDEEEEDVKLLSISGKRSAPGGGSKVPQK<br>KVKLAADEDDDDDDEEDDDEDDDDDDFDDEEAEEKAPV<br>KKSIRDTPAKNAQKSNQNGKDSKPSSTPRSKGQESFKK<br>QEKTPKTPKGPSSVEDIKAKMQASIEKGGSLPKVEAKFIN<br>YVKNCFRMTDQEAIQDLWQWRKSL |
| Wild-type SURF6, in vitro | MGSSHHHHHHSQDLENLYFQGSMASLLAKDAYLQSLAK<br>KICSHSAPEQQARTRAGKTQGSETAGPPKKKRKKTQKKF<br>RKREEKAAEHKAKSLGEKSPAASGARRPEAAKEEAAWA<br>SSSAGNPADGLATEPESVFALDVLRQRLHEKIQEARGQG<br>SAKELSPAALEKRRRRKQERDRKKRKRKELRAKEKARKA<br>EEATEAQEVVEATPEGACTEPREPPGLIFNKVEVSEDEP<br>ASKAQRRKEKRQRVKGNLTPLTGRNYRQLLERLQARQS<br>RLDELRGQDEGKAQELEAKMKWTNLLYKAEGVKIRDDE<br>RLLQEALKRKEKRRAQRQRRWEKRTAGVVEKMQQRQD<br>RRRQNLRRKKAARAERRLLRARKKGRILPQDLERAGLV |

**Table S3: Primer sequence of SURF6 (C19S) construct**.

| Construct Name | Primer Sequence |
|---|---|
| SURF6 (C19S), in vitro | Forward: 5’ CAA TCG TTA GCG AAG AAA ATT TCT TCA CAT AGT GCC CCA GAG 3’<br>Reverse: 5’ CTC TGG GGC ACT ATG TGA AGA AAT TTT CTT CGC TAA CGA TTG 3’ |
